# Are human super-predators always super-scary? A meta-analysis of wild animal behavioural responses to human interactions

**DOI:** 10.1101/2024.08.27.609826

**Authors:** Shawn Dsouza, Kartik Shanker, Maria Thaker

**Affiliations:** Centre for Ecological Sciences, Indian Institute of Science, Bengaluru, India

**Keywords:** risk disturbance, nonconsumptive, predator – prey interactions, fear, foraging, vigilance, movement

## Abstract

Humans interact with wild animals through lethal and non-lethal activities. While theory predicts that these interactions should alter animal behaviour, the relative magnitude of impact is not well understood. We conducted a systematic review and meta-analysis of the past 30 years of research, focusing on changes in foraging, vigilance, and movement behaviours in wild animals. We found that lethal human activities (e.g., hunting) cause significant behavioural changes, with targeted species increasing vigilance and reducing foraging in affected areas. Active non-lethal activities (e.g., tourism) elicited similar but weaker patterns, with many species showing little to no change in their behavioural responses. In contrast, passive non-lethal interactions (e.g., roads) produced highly variable responses. Overall, human-induced fear elicits responses in wild animals that broadly align with predictions from the risk allocation hypothesis. However, the magnitude and direction of animal responses depend on the type of human activity and the ecological context. The most pronounced behavioural changes occurred where humans posed a direct lethal threat. Gaps in the literature, uneven data within and across species, and limited information on the history or context of interactions currently limit our ability to better predict when and why animals change their behaviour in response to humans.

## Introduction

Humans have impacted natural environments in profound and far-reaching ways. This influence stems partly from their unique ecology, which has enabled humans to occupy every ecoregion of the world and play multiple functional roles (Darimont et al. 2023). For the majority of their evolutionary history, humans have hunted, trapped, and fished other animals (Treves & Naughton-Treves 1999). Social organisation and use of tools have enabled humans to efficiently target wild animals across multiple trophic levels simultaneously (Treves & Naughton-Treves, 1999). Thus, as predators, humans are more deadly than most other predators (Darimont et al. 2015). In fact, the mortality rates of prey species from human predators, such as hunters and fishers, are significantly higher in magnitude than the sum of kills by all natural predators in both terrestrial and marine ecosystems (Darimont et al. 2015). Humans have therefore been referred to as generalist super-predators (Clinchy et al. 2016) and they create truly risky conditions for animals that they intentionally kill in the wild (Oriol-Cotterill et al. 2015).

In addition to hunting and fishing, humans also interact with wild animals through non–lethal activities (Boyle & Samson, 1985), which can be further distinguished as either active or passive. Active non-lethal interactions include tourism, walking in parks, hiking in nature reserves, scuba diving, and snorkelling. Studies have shown that these activities can have a negative impact on animals, even when humans in their habitats are not actively killing them (e.g., Brown et al. 2012; Loehr et al. 2005; Montero-Quintana et al. 2020; Newey 2007). Passive non-lethal interactions are even more pervasive, as they include engineered alterations to environments. Roads and settlements, in particular, have the greatest potential to influence the behaviour of animals (e.g., Cappa et al. 2017; Mehlhoop et al. 2022, etc.). Some types of "passive" human interactions may also cause significant mortality, such as through collisions on glass (Loss et al. 2015) and the high occurrence of road-kill globally (Benson et al. 2015). Although some studies suggest that non-lethal human activities can negatively influence animal behaviours (Boyle & Samson 1985; selected examples: Papouchis et al. 2001; Davis et al. 1997; Westekemper et al. 2018; Ladle et al. 2019; Nevin & Gilbert 2005), other studies show that animals habituate quickly to human encounters and may even benefit from these encounters (see Smith et al. 2024, Gaynor et al. 2025). As such, non-lethal interactions with humans can have either positive or negative effects on wild animals (see Smith et al., 2024).

Given that humans interact with animals in the wild with both lethal and non-lethal intent, the individual and combined effects of these interactions on animal behaviours can potentially be high. Decades of research have shown that animals balance the competing demands of energy gain and safety under the risk from natural predators (Sih, 1984; Lima & Dill, 1990; Brown & Kotler, 2004). At the most fundamental level, prey are expected to adjust their vigilance and foraging behaviours in response to both the presence of predators and the degree of risk they pose (‘risk allocation hypothesis’; Lima & Bednekoff, 1999). This expectation has been extended to predict wild animal responses to humans. Previous studies have suggested that animals may perceive humans (Clinchy et al., 2016) and their alterations to the environment (Frid & Dill, 2002) as risky, with non-lethal interactions inducing a fear response that is almost as high as that caused by lethal interactions (Clinchy et al., 2016; Frid & Dill, 2002). In a recent narrative review, Smith et al. (2024) outlined a framework that describes the pathways by which lethal and non-lethal interactions with humans can alter the behavioural and physiological responses of animals, with consequences for population growth and abundance. They conclude that the perceived risk of humans may induce phenotypic changes in wild animals, but, as expected, the evidence is mixed (Smith et al., 2024). What is still missing is a quantitative assessment of the relative magnitude of effect for different types of human activities, which would illustrate when, why, and how wild animals respond to human-induced risk. In particular, it remains unclear whether non-lethal interactions between humans and animals are as strong as lethal interactions in eliciting fear and changing behaviours.

In this paper, we synthesise and review the current evidence for wild animal behavioural responses to the risk of humans in their environment. We categorised studies based on whether wild animals were exposed to interactions with humans of lethal intent (hunting, fishing), active and non-lethal intent (ecotourism, hiking), and passive and non-lethal intent (roads and human settlements) (Figures 1 and 2). We focused on long–term changes in foraging behaviours (bite rates and foraging time), vigilance (vigilance time and proportion), and movement (displacement, home range size, and movement rate) of animals in response to human interactions. Immediate responses, such as flight or escape, were excluded from this meta-analysis as they occur at short time scales and may not have long-term fitness consequences in and of themselves. Physiological responses, such as changes in hormone levels (Sherrif et al., 2020; Smith et al., 2024), were also not included here, as these physiological changes can influence multiple behavioural pathways and are variable across taxa (Sheriff and Thaler, 2014).

**Figure 1:**
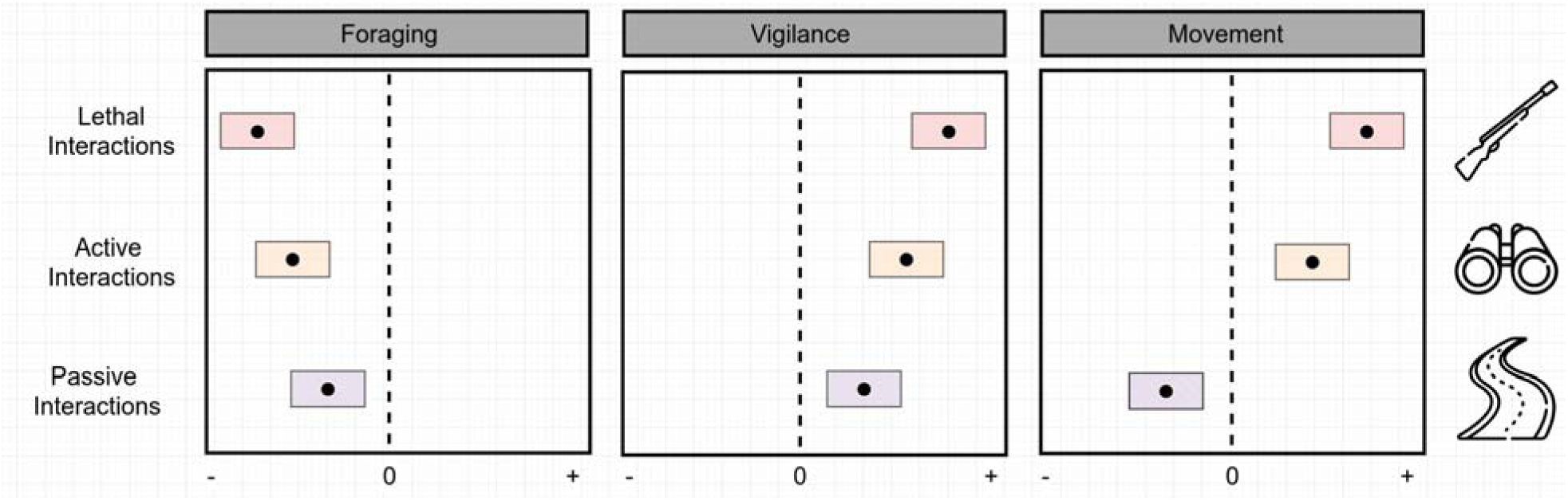
The different scenarios in which humans interact with wild animals and the hypothesised consequences on animal behaviour. We expect a gradient of response, with the most substantial behavioural changes in animals in areas where interactions with humans are of lethal intent (e.g., hunting and fishing), followed by non-lethal but active interactions (e.g., ecotourism), and non-lethal but passive interactions (e.g., roads). Values below 0 represent a reduction in the behaviour, while those above 0 represent an increase relative to animals in area without that human activity.

**Figure 2:**
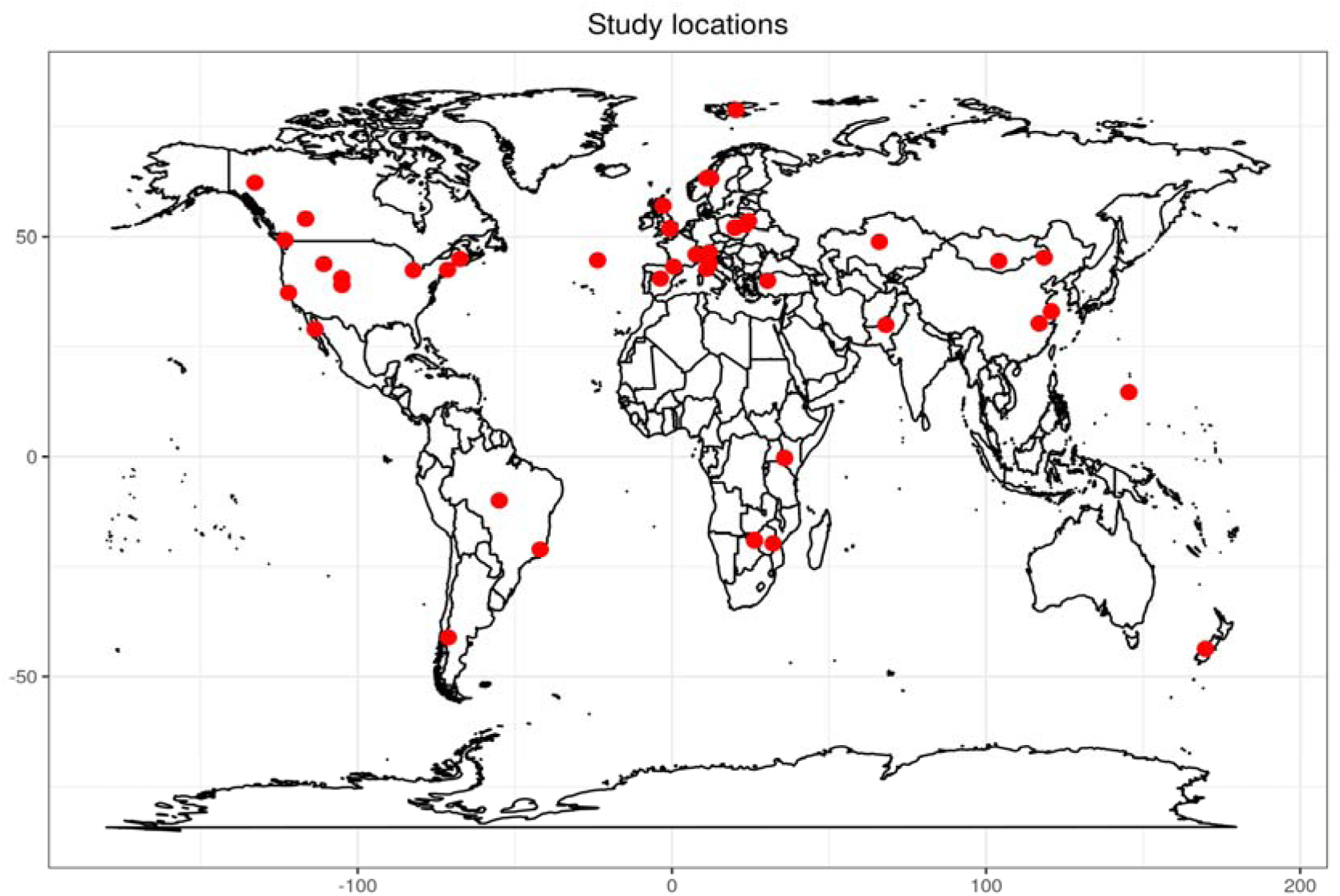
Geographical distribution of studies included in the systematic review.

From the systematic review of published studies, we derived effect sizes that describe the magnitude of behavioural change across these categories of human interactions. We then conducted a meta-analysis to determine whether active or passive non-lethal interactions with humans influence wild animal behaviours to the same extent as lethal human interactions. We predict that, in areas where humans have lethal intent towards animals, these animals will strongly invest in anti-predator behavioural strategies, spending more time being vigilant, less time foraging, and increasing their movement rate or displacement (Figure 1). We expect a lower intensity of anti-predator responses to active and passive non-lethal interactions with humans, as wild animals may perceive such interactions as non-threatening or even beneficial based on cues given off by humans in the habitat (Wirsing et al. 2021, Figure 1). We also tested the potential influence of body size and trophic level on the magnitude of behavioural effects based on the hypothesis that these factors influence which animals are targeted by the lethal actions of humans (i.e., they are hunted, trapped, or fished). Finally, we highlight where gaps in the literature have yielded skewed conclusions about human impacts on the behaviour of wild animals and suggest avenues for future research in this field.

## Methods

To determine the strength of behavioural change (i.e. effect sizes) caused by human activities, we conducted a systematic literature survey that included studies where foraging, vigilance or movement behaviours were measured in wild animals in response to potentially lethal human interactions (hunting, including poaching and trapping, and fishing), non-lethal but active interactions (walking in parks, hiking in nature reserves, off-road bicycling and animal watching), and non-lethal but passive interactions (roads and human settlements) (Figure 2). We focus on only wild animal behaviour that is indicative of long-term foraging safety trade-offs (Lima & Dill, 1990); thus, we excluded studies of escape behaviour (typically measured as flight initiation distance) and studies that measure animal physiology. We also excluded studies where human interactions were induced to simulate a natural predator approach, if there was no information about the underlying context of human interaction (i.e., lethal and non-lethal intent).

### Search and literature database

We conducted a scoping search using the Web of Science database and its advanced search function. Our initial search term was "(risk OR fear) AND (human OR anthropogenic) AND (behavio*) AND ALL=(predat*)". The search terms were applied across all fields. The initial search returned 2,319 results on January 12, 2023. We downloaded the data and conducted a literature mapping analysis to identify additional search terms and updated our search string to "(human OR anthropo*) AND (risk OR fear OR NCE OR "trait mediated effect" OR nonconsumptive* effect) AND (behavio*) AND (predat*)". Using our updated search string, we conducted searches on the SCOPUS and WOS databases, as well as OpenThesis, for grey literature on February 14, 2023. We then used Rayyan, a literature management and screening software, to deduplicate our database (Foo et al., 2021; Ouzzani et al., 2016), which resulted in 7,562 abstracts. This review was not registered.

### Screening

We piloted our initial screening protocol on 100 randomly selected papers using Rayyan (Ouzzani et al. 2016). We determined the inclusion and exclusion criteria by evaluating titles, abstracts, and keywords to identify the population studied, the exposure or intervention applied to the population, the control (if any), and the outcome (see Appendix A for detailed inclusion criteria). The initial screening was conducted by three independent researchers, who discussed conflicting decisions on inclusion or exclusion of abstracts until a consensus was reached. We applied the finalised screening protocol to all abstracts in our database and were left with 436 papers to evaluate further. Papers chosen for inclusion during our initial screening were full-text screened. We included 71 studies at this stage (see Appendix A for our complete screening protocol).

### Backward and forward search

We identified key studies and reviews during our screening for backward and forward search. The key studies (n = 14) included major reviews, studies that first addressed the effects of human interactions on animal behaviour, and concept papers (see Appendix B for a complete list). Backward search identified papers cited by these studies and reviews, and forward search identified papers that cited these key papers. We used a citation chaining tool called ResearchRabbit to execute the search on March 12, 2024. We added the results to our existing database and deduplicated it to create the final paper database for the review (Foo et al. 2021). After screening the additional abstracts, we screened the full texts of 40 papers, and 14 were added. Our final sample included 44 studies for our meta-analysis and an additional 41 for review (See Appendix B for a complete list). We have included a PRISMA (EcoEvo) flowchart and checklist in Appendices A and F, respectively (O’Dea et al., 2021; Page et al., 2021).

### Data collection

We collected data on the behavioural responses, study species, trophic level (Consumer, Primary Predator, and Secondary Predator), location of the study, treatments, controls, and sample size. Trophic level, dietary guild, and body mass of the study species from each study were curated from Mammalbase (Lintulaakso, 2021), the Animaltraits database (Herberstein et al., 2022), the Handbook of the Birds of the World (Del Hoyo et al., 1992), and Fishbase (Froese & Pauly, 2010). We included omnivores as either Primary or Secondary predators based on their diets (e.g., bear species were included as secondary predators, see Appendix A).

For the animal responses, we extracted data on the mean effects of foraging, vigilance, and movement behaviours, along with associated variance measures (Figure 3). Studies that quantify foraging and vigilance behaviours used focal individual sampling and scan sampling. The measured behaviours were time spent vigilant, time spent foraging, frequency of vigilance behaviour in a group, rate of individual vigilance behaviour, and foraging (bite) rates. We included movement studies that measured movement rates (speed), displacement (distance travelled), and home range size using GPS tags, collars, and transmitters. Home range size was included as it is a function of displacement and speed in a given sampling period. Where results were not clearly reported, we either used raw data to calculate mean effects and variance measures or used a plot digitiser to extract data from figures when raw data were not available. Studies in which data extraction was not possible were excluded from our meta-analysis and were used only for qualitative review. We excluded papers that only reported test statistics from our meta-analysis (Mikolajewicz & Komarova, 2019; see Appendix B for a complete list of studies).

**Figure 3:**
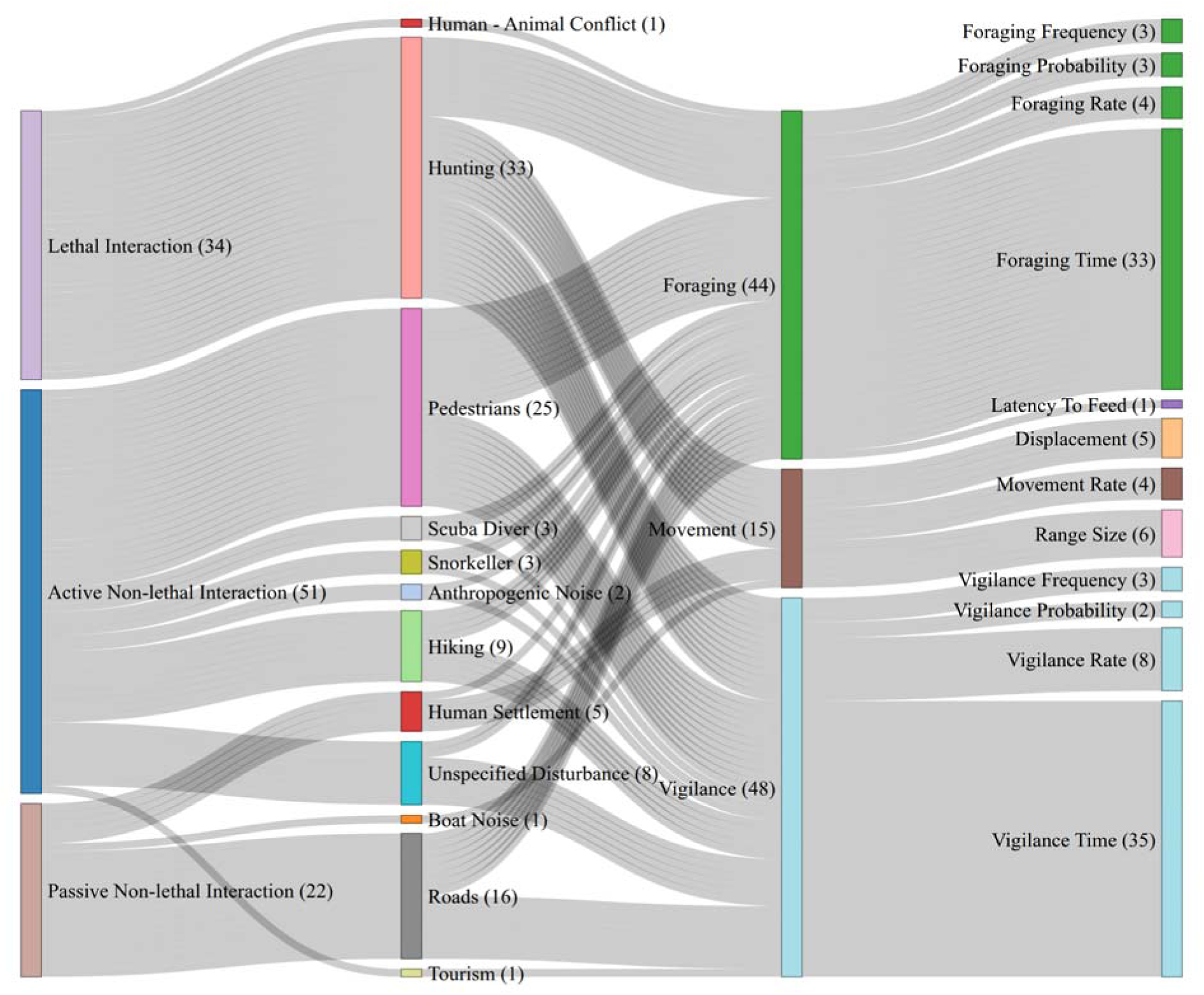
Distribution of studies included in the meta-analysis. Shown here are the various hu- man activities that were considered under lethal, active non-lethal, and passive non-lethal, as well as the specific behavioural measurements of wild animals within the broad categories of foraging, vigilance, and movement. The number of studies at each node is indicated in brackets.

Several types of study approaches were used to measure animal responses to human activities. The majority of studies were observational or natural experiments. Spatial contrasts included designs in which behaviours were compared across protected and unprotected areas for lethal interactions (n = 8), areas with and without tourists for active interactions (n = 6), and areas close and far away from roads or settlements for passive interactions (n = 5). Studies also employed auditory stimuli (n = 5), such as road noise or human speech, in areas with hunting pressure or high traffic to test for the effect of lethal interactions and passive interactions, respectively. Other experimental studies employed direct human disturbance to simulate active interactions (n = 6) and lethal interactions in areas with hunting (n = 3). Four studies utilized temporal contrast in human activities, such as hunting closures and tourist seasons, to investigate the effects of active (n = 1) and lethal (n = 3) interactions. Finally, ten studies did not implement or utilize a contrast for their comparison. Instead, they determined correlations between human activities and distance from the road for both active and passive interactions.

### Analysis

We divided the studies into categories based on their study designs as experimental or observational. We converted proportional and percentage data (means and measures of uncertainty) to absolute values (number of individuals or time in seconds). We converted all measures of uncertainty to variance (using sample size to convert SE to SD) before analysis.

Furthermore, we calculated standardised mean difference for studies that reported an explicit treatment and control (negative or positive), as 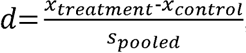 where is the mean of treatment and control outcomes respectively, and 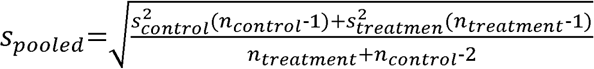 here is the sample variance and is the sample size.

All the correlational studies reported unstandardised regression coefficients (*b*), and thus, we calculated the standardized mean differences from *b* as 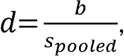 where 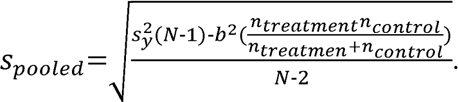 Here, *N* is the total sample size (Lipsey and Wilson, -2001). Where generalized linear models were used or where data were transformed prior to fitting a linear model, we back-transformed the coefficients to the original scale. We used a control-treatment contrast, with the treatment being those groups that were exposed to lethal or non-lethal interactions, and the control being groups not exposed to interactions. The model coefficients of studies that used a control:treatment contrast were multiplied by -1 to maintain consistency across studies. We assigned each study a unique identifier (study ID). When extracting multiple data points from studies, we assigned a unique identifier to each data point (data ID).

We first fit an intercept-only multilevel meta-analytic (MLMA) model using the metafor package (Viechtbauer 2010) with species, study, and data ID as random effects to determine the degree of heterogeneity across studies after accounting for known sources of variation as follows:

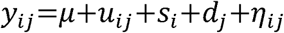

Where, is the effect from *j^th^* data point of the *i^th^* study, μ is the overall mean across studies, is the *i^th^* estimate within the *j^th^* study, is the random effect due to the *i^th^* study, is the random effect due to the *j^th^* data point, and is the error of the *j^th^* estimate from the *i^th^* study. The random effect framework assumes that each study has its own proper effect size, which is derived from a population of true effect sizes. Since this meta-analysis involves a comparison across taxa in different habitats and geographies, the random-effects framework is appropriate (Nakagawa et al., 2022; Noble et al., 2022). We utilized the Open Tree of Life API via the ℈rotl’ package to construct a phylogenetic tree of species included in our meta-analysis (Michonneau et al., 2016; OpenTreeOfLife et al., 2019). We also fit a phylogenetic model with relatedness as a random effect.

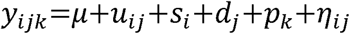

where is a random effect due to the *k^th^* species. Phylogeny did not capture significant variation across all outcomes (Chamberlain et al. 2012; Cinar et al. 2022). We thus used a model with only study identity and data identity as random effects. We used the *Q* statistic to determine leftover variation. Furthermore, we used both and to determine heterogeneity across studies (Harrer et al. 2021). We used the final MLMA to determine whether there was a significant overall effect across studies. Finally, we used the MLMA to perform a multiple regression to determine the effect of the type of human interactions and animal size on the observed effects. We have described our analysis of sensitivity, robustness and bias of our dataset in Appendix D. We report the mean and 95% confidence intervals of the strength of each behavioural response we have considered in our meta-analysis; foraging, vigilance, and movement, to each type of human interaction we have considered in our meta-analysis; lethal, active non-lethal and passive non-lethal, as measured by standardised mean difference of responses to treatment and responses to control. Additionally, we present regression analyses examining the effect of different types of human interaction on these behaviours, including T-statistics and significance levels.

All data preparation and analysis were conducted using R 4.0.1 (R Core Team, 2024). Data and code are available at Dsouza *et al*. (2025).

## Results

After screening, we included 85 studies in our systematic review, a majority of which were published in the past two decades (Appendix A). Of these, 44 studies provided sufficient information to be included in the meta-analysis, encompassing 38 species. These studies were widely distributed across geographies and habitats, ranging from 78 ° N to 43 ° S and 169 ° E to 123 ° W (Figure 2). Across outcomes, 24 studies reported measures of foraging behaviours, which included bite rate and time spent foraging, 29 studies focused on vigilance behaviours, which included time spent vigilant and vigilance rate, and nine studies measured movement behaviour, which included home range size, displacement, and rate of movement (Figure 3). Eighteen studies reported data from more than one major category of behaviour. The majority of these papers (n = 36) studied primary consumers, such as *Cervus elaphus* and *Capreolus capreolus* (Figure 4, Appendix A). Only seven studies included secondary and tertiary consumers (Figure 4); thus, statistical comparisons across trophic levels were not conducted.

**Figure 4:**
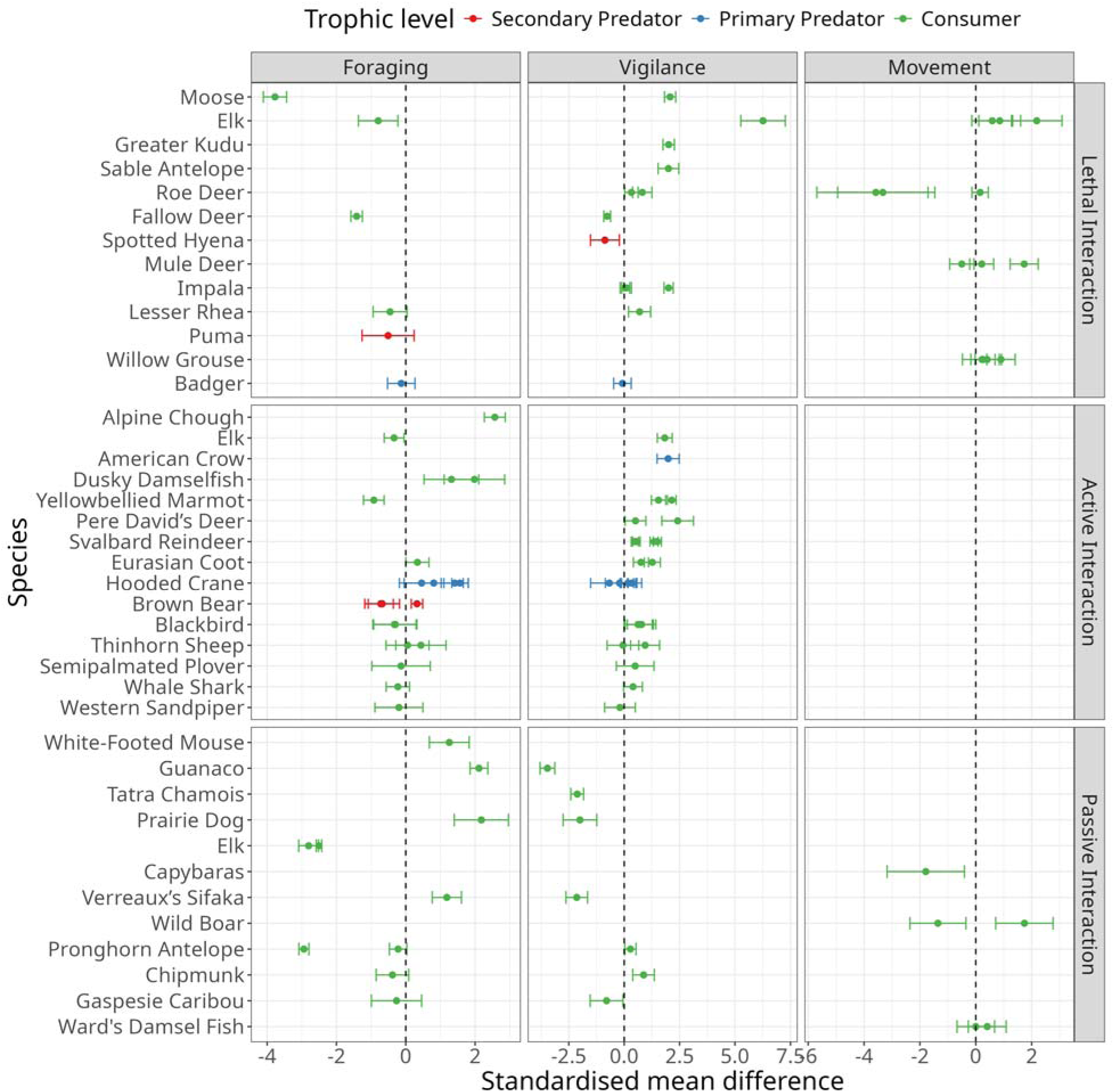
The effect of human interactions (lethal, active non-lethal, passive non-lethal) across taxa and trophic levels. Negative values on the x-axis indicate a reduction in that behaviour. In contrast, positive values indicate an increase in that behaviour, relative to control conditions where no human activity is present.

### Changes in behaviours across types of human interactions

Across studies, the overall effect of human interactions on foraging behaviours was adverse (SMD = -0.178, 95% CI = -0.738 – 0.382, = -0.624, p = 0.533) and on vigilance was positive (SMD = 0.641, 95% CI = 0.292 – 0.991, = 3.593, p = 0.0003). However, the effect of human interactions on movement behaviour across studies was negligible (SMD = 0.191, 95% CI = - 0.219 – 0.594, = -0.172, p = 0.868, Appendix E). Upon fitting the MLMA model, significant heterogeneity remained among studies that was not accounted for by random effects (Appendix E).

Foraging behaviour of animals decreased significantly in response to lethal human interactions (SMD = -1.193, 95% CI = -2.279 – -0.108, = -2.154, p = 0.031), and to a lesser extent in response to passive non-lethal interactions (SMD = -0.255, 95% CI = -1.392 – 0.882, -0.439, p = 0.66, Figure 4). Active non-lethal interactions had an overall significant positive effect on foraging (SMD = 0.331, 95% CI = -0.115 – 0.778, = 1.455, p = 0.145, Figure 4). Vigilance behaviour increased significantly in response to lethal interactions (SMD = 1.1, 95% CI = 0.014–2.186, = 2.991, p = 0.003) and active non-lethal interactions (SMD = 0.829, 95% CI = 0.549– 1.11, = 5.798, p = 0). However, vigilance behaviour was significantly reduced in passive non-lethal interactions (SMD = -1.339, 95% CI -2.042 – -0.054, = -2.042, p = 0.041). There was no significant overall effect of human interactions on movement (Figure 5).

**Figure 5:**
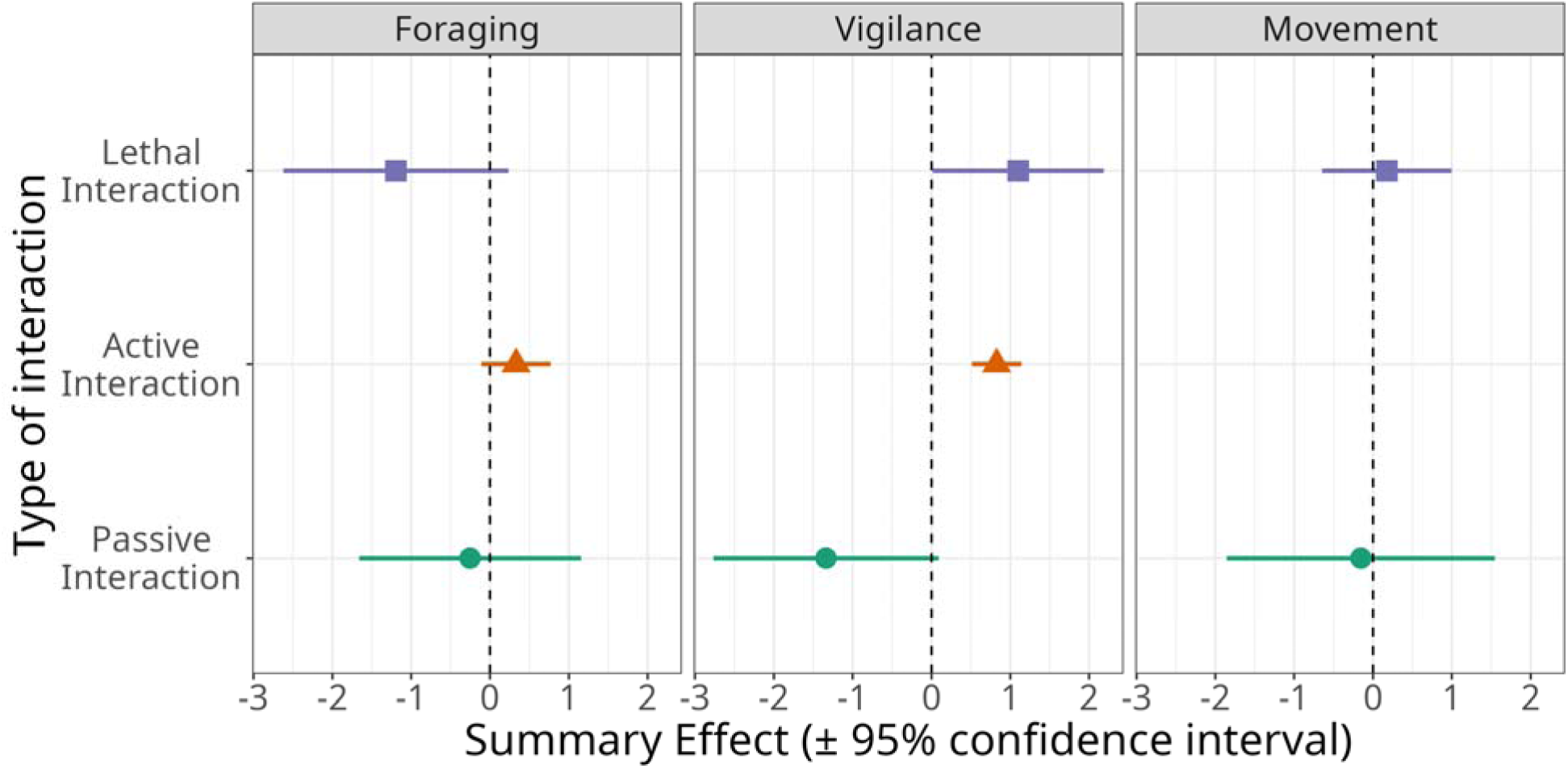
Summary effect of human interactions on foraging, movement, and vigilance behaviour across studies included in this meta-analysis. For animal movement, active non-lethal interactions were excluded due to the low sample size.

Overall, when considering all studies in this analysis, foraging behaviours were significantly more suppressed by lethal human activities than by active non-lethal human interactions (β = - 0.955 ± 0.51, *T_34_* = -1.874, p = 0.07, Figure 5). There was no significant difference in the magnitude of effect on foraging behaviours between active and passive non-lethal human interactions. The magnitude of behavioural effects on vigilance behaviour due to lethal human interactions was significantly higher than that of active non-lethal interactions (β = 1.347 ± 0.553, *T_46_* = 2.433, p = 0.019, Figure 5). There were no significant differences in the effect of human interactions on movement behaviours across the three types of interactions (Appendix E, Figure 5).

The body size of the animal did not influence the effect of human activities on foraging (β = 0 ± 0, *T_31_* = -0.31, *p* = 0.75), movement (β = 0.005 ± 0.006, *T_16_*= 0.948, *p* = 0.37), or vigilance behaviour (β = 0 ± 0, *T_41_*= 0.089, *p* = 0.92).

## Discussion

The term human “super-predator" was coined to highlight the efficiency, lethality, and scale at which human predators operate (Clinchy et al. 2016; Darimont et al. 2015). Recent frameworks, such as those proposed by Smith et al. (2024), have begun to assess the ecological impacts of humans as super-predators. Building on this, our study offers a first step toward quantifying these effects by conducting a meta-analysis of wild animal behavioural responses to human interactions, differentiating between those with lethal and non-lethal intent. Our analysis, spanning a broad geographic and taxonomic range (Figure 2), reveals that lethal interactions exert the most decisive influence on overall wild animal behaviour, followed by active non-lethal interactions. In contrast, passive interactions, such as those with roads and human developments, had a suppressive effect on wild animal vigilance behaviours but did not show consistent effects on foraging or movement (Figure 5). These results broadly align with the risk allocation hypothesis, wherein prey respond most strongly to predators when they are lethal and active (Lima & Bednekoff 1999; Ferrari et al. 2009). Importantly, we found significant variation in wild animal responses to human activity across different interaction types (Figure 4), underscoring the strong context dependence of these effects across species. Below, we discuss the findings of our meta-analysis within the broader context of human-animal interactions.

### Are humans always super-scary when present?

Darimont et al.’s (2015) analysis revealed that humans have killed a larger biomass of prey species than most other predators in their respective ecosystems. Given the efficiency and scale at which humans kill animals, potential prey should respond strongly to human hunters (Clinchy et al. 2016; Crawford et al. 2022). Our meta-analysis revealed that, across taxa and geographical locations, wild animals exhibit significant alterations in both their foraging and vigilance behaviours in areas where lethal interactions with humans occur. In line with the foraging– vigilance trade-off (Brown & Kotler, 2004), we found that animals reduced foraging and significantly increased vigilance in response to human predators, similar to their response to other predators in their environment.

There was, however, some variability in the strength of animal responses to humans with lethal intent (Figure 4). Moose (*Alces alces*), elk (*Cervus elaphus*), and fallow deer (*Dama dama*) exhibited the most substantial reductions in foraging behaviour., while elk showed the strongest increase in vigilance. In contrast, roe deer (*Capreolus capreolus*) displayed the most substantial reduction in movement. The magnitude of behavioural change in impala (*Aepyceros melampus*) depended on the study. Matson et al. (2005) recorded much lower vigilance for impala than Crosmary et al. (2011), who measured animals approaching water—a context where need and risk intersect. This pattern highlights how, for some species, internal state modulates risk sensitivity, as supported by broader research on state-dependent anti-predator behaviour. (Wirsing et al., 2021; Zanette et al., 2020).

Similar to lethal interactions with humans, non-lethal interactions with humans also elicited an overall positive effect on wild animal vigilance behaviour. (Figure 5). The aggregate effect also indicated an increase in foraging in the presence of non-lethal humans; however, this effect was neither large nor significant (Figure 5). According to the risk–disturbance hypothesis, animals are expected to perceive even non-lethal human disturbance similarly to the threat of predation (Frid & Dill 2002). Overall, we found that wild animals responded to non–lethal active human interactions to a lesser extent than lethal disturbance (Figure 5). Variation in the strength of response to non-lethal active interactions was also lower than that seen for lethal interactions, indicating a more consistently muted response to human presence across taxa.

### Human-altered environments have variable effects

Over the past two centuries, the pace and scale of human transformation of ecosystems have accelerated dramatically, leading to rates of change unparalleled in the planet’s history (Estes et al. 2011; Sih 2013). We found that passive human interactions, such as roads and human settlements, did not significantly influence the foraging or movement behaviour. of wild animals.

However, vigilance behaviours were significantly lower in response to human-altered environments (Figure 5). While this may, in part, reflect the paucity of studies that directly assess these effects, we nevertheless found substantial variation in behavioural responses to passive human interactions across studies (Figure 4). Unlike active lethal interactions, which typically produced reduced or non-significant effects, responses to passive interactions were often large in magnitude and highly variable.

Some wild animals appear to benefit from passive human interactions. The human–shield hypothesis proposes that human presence can alter interspecific interactions by deterring predators or dominant competitors and creating safe areas for prey or subordinate competitors (sensu Berger 2007, Moller 2008; Proudman et al. 2021; Shannon et al. 2014; Abrahms et al. 2025). Roads, for example, typically remove cover that is typically used by ambush predators, paradoxically making areas adjacent to roads "safer" for specific prey species (Berger 2007; Moll et al. 2017). From our meta-analysis, evidence of a positive effect of passive human interactions, predicted by the human-shield effect (Berger 2007), can be seen in the increased foraging and reduced vigilance behaviours of prairie dogs (*Cynomys ludovicianus*), white-footed mice (*Peromyscus leucopus*), and Verreaux’s sifakas (*Propithecus verreauxi*) (Shannon et al. 2014; Giordano et al. 2022; Chen-Kraus et al. 2022; Figure 4). It should be noted, however, that roads can also serve as ecological traps, offering new foraging opportunities while increasing mortality risks (Smith et al. 2021).

Human-altered environments may not always confer advantages. In some cases, human development increases predation risk by opening habitats and making it more difficult for prey to minimise detection and escape (Smith et al. 2016). Human infrastructure also poses direct threats to wildlife via vehicle collisions, poisoning, or conflict with livestock interests (Harris et al. 2018; van der Kolk et al. 2024; Loss et al. 2015). Consistent with these risks, several studies have documented reductions in foraging and heightened vigilance of elk (*Cervus elaphus*) near roads and other passive disturbances (Ciuti et al. 2012; Shannon et al. 2014; Jayakody et al. 2008). Foraging responses in pronghorn (*Antilocapra americana*) were mixed, potentially attributable to differences in study design. For instance, Gavin et al. (2006) employed distance from roads as a continuous control variable, while Shannon et al. (2014) used categorical comparisons between disturbed and undisturbed road sites. Ultimately, whether animals perceive human-altered environments as refuges or threats varies significantly by species and ecological context (Wirsing et al., 2021; Smith et al., 2021).

### Comparing the effects of human interactions within species and individuals

The biggest challenge in understanding the relative impacts of lethal and non-lethal human interactions on wild animals is that only three studies have tested these effects in the same species. An exceptionally informative example from our meta-analysis is elk (*Cervus elaphus*), the only species in our dataset for which foraging and vigilance behaviours were assessed under both lethal and non-lethal human disturbance conditions (Appendix A: Table A1; Figure 4). Elk consistently reduced foraging and increased vigilance, regardless of interaction type, and in some contexts, displayed a pronounced increase in vigilance even without a concomitant change in foraging (Jayakody et al. 2008). Additionally, lethal interactions with humans induced a marked increase in elk movement rates (Ciuti et al. 2012; Shannon et al. 2014). Such strong anti-predator behavioural responses towards humans, regardless of whether they have lethal or non-lethal intent, may reflect a long history of hunting pressure on elk (Ciuti et al. 2012).

In addition to elk, several other species in our dataset—including willow grouse, brown bear, whale shark, roe deer, fallow deer, and badgers—have documented histories of lethal or antagonistic interactions with humans (See appendix A: Table A1). However, the nature and strength of their behavioural responses were highly variable (Figure 4). These examples underscore that a mere history of lethal encounters is insufficient to predict how animals will respond to humans in their environment. The presence and magnitude of behavioural responses to humans may depend on several additional factors, such as the intensity of hunting or persecution, the nature of human activities in the habitat, the duration since the cessation of hunting or persecution, and current forms of human interaction (Smith et al., 2021, 2024).

The criteria of our meta-analysis precluded us from including some other interesting examples of when lethal and non-lethal interactions with humans combine to influence animal behaviour. For example, moose (*Alces alces*) in Norway avoid human infrastructure in hunting areas, suggesting a synergistic effect of different types of human disturbance (Mehlhoop et al. 2002). Such changes in behaviour may have compounding adverse effects on wild animals. Roe deer and wild boar alter their habitat use in response to hunting pressure, which in turn makes them more susceptible to death from vehicle collisions (Zuberogoitia et al. 2014). These multiple-exposure scenarios highlight a critical area for future research that our meta-analysis was unable to address fully.

Approximately half of the studies in our meta-analysis examined the effects of human interactions on both foraging and vigilance behaviours of individuals (n = 18). This allows us to comment on direct foraging–vigilance trade-offs (Brown and Kotler, 1991). Qualitatively, there is reasonable consensus among these studies that animals reduce foraging and proportionally increase vigilance in response to both lethal and active non-lethal interactions with humans. Similarly, there is consensus that animals increase foraging and reduce vigilance in response to human-altered habitats (Appendix A: Table A3).

While our meta-analytic approach enables unitless comparison across studies (Noble et al., 2022), this does not alter the fact that different behavioral metrics may be indicative of different underlying processes, and different study designs may yield different conclusions. For example, movement and space use in response to passive human disturbances are governed by distinct behavioural processes. Wild boar (*Sus scrofa*) in urbanised environments occupy smaller home ranges than those in undisturbed forests, yet move greater distances within these ranges, reflecting both reduced spatial requirements and increased activity (Podgorski et al. 2013; Zanette et al. 2020). Another notable example is the contrasting movement responses of roe deer (*Capreolus capreolus*) to the presence of lethal human activity (Figure 4). However, this discrepancy is plausibly attributable to methodological differences between studies. Specifically, Grignolio et al. 2011 used a temporal control to test the effect of hunting, whereas Picardi et al. (2009) employed a spatial control. These mixed responses indicate that different behavioral metrics and designs can result in varied responses to human disturbance within the same species.

### Perception of risk and the role of habituation

Contrary to expectation, some animals showed little change in behaviour in response to lethal interactions with humans. Despite being hunted, puma (*Puma concolor*, Smith et al. 2017), badgers (*Meles meles*, Clinchy et al. 2016), and impala (*Aepyceros melampus*, Matson et al. 2005), which were included in our meta-analysis, did not change either foraging or vigilance behaviour or both (Figure 4). These species may be under-reacting to humans in their environment, and their lack of response may reflect the nature of how these animals are killed and their ability to identify and assess human predator cues (Smith et al. 2021). For example, animals may not respond the same way to humans with guns and traps as they provide different cues. On the other hand, some animals seem to overreact, such as those that significantly change their behaviours in response to non-lethal human interactions, leading potentially to high energy investment or chronic stress (e.g., Blumstein 2003; Westekemper et al. 2018; Larson et al. 2016; Nunes et al. 2019 review: Larson et al. 2016).

Animals may also habituate to human encounters or human-induced changes in the environment, which may explain the subdued or non-significant responses to active non-lethal human interac- tions. In our meta-analysis, blackbirds (*Turdus merula*), thinhorn sheep (*Ovis dalli*), whale sharks (*Rhincodon typus*), semipalmated plovers (C*haradrius semipalmatus*), and western sand- pipers (*Calidris mauri)* showed no change in either foraging or vigilance in response to active human activity in their environment (Fernandez-Juricic et al., 2001; Loehr et al., 2005; Montero- quintana et al., 2020; Murchison et al., 2016, respectively). Developmental stage may also medi- ate responsiveness; for instance, while adult thinhorn sheep increase vigilance in response to non-lethal disturbance, juveniles do not, and only juveniles show flexibility in their foraging behaviour. (Loehr et al. 2005). Similar patterns occur in hooded cranes (*Grus monacha*), which vary vigilance but not foraging responses across age classes (Li et al. 2015). These observations suggest habituation, wherein repeated exposure to non-lethal humans results in a dampening of anti-predator responses over time.

A recent meta-analysis of camera trapping data from 21 countries during the COVID-19 pandemic also suggests that habituation may explain significant variation in animal responses to non-lethal human presence (Burton et al., 2024). For example, Uchida & Blumstein (2021) found that yellow-bellied marmots (*Marmota flaviventer*) in highly disturbed forests grew habituated to humans after repeated exposure and reduced their flight initiation distances. Similar examples have been cited from tourist interactions with animals in the African savannah (Knight, 2009), interactions between killer whales and Hector’s dolphins with whale watchers (Bejder et al., 1999; Williams et al., 2002), and interactions between researchers and penguins (Shelton et al., 2004).

As with non-lethal active human interactions, not all taxa included in this meta-analysis responded to passive human interactions: chipmunks (*Tamias striatus*) and Gaspésie caribou (*Rangifer tarandus caribou*), for example, show limited behavioural shifts to passive human presence for reasons that are unclear (Giordano et al. 2022; Lesmerises et al., 2017). The heterogeneity in risk perception and habituation contributes to the overall variation observed in behavioral responses across taxa, highlighting the need for further research encompassing a broader diversity of species and behavioral metrics (Figure 1).

### Context-dependent variability in behavioural responses to humans

The results of our meta-analysis reveal the importance of context in how animals respond to humans. Ecological and biological factors, such as trophic level, body size, and group size, can influence the perception of risk by animals. Unfortunately, there were too few studies in the meta-analysis to enable a statistical evaluation of these effects. These, however, are important enough to warrant some discussion.

When humans kill top predators, they create a new risk that these animals have not previously experienced. Predators may or may not be equipped to respond to new threats, leading to many extirpations (e.g., wolves in Yellowstone (Laundré et al. 2001), bears in Alaska (Merkle et al. 2013), puma in California and Florida (Nickel et al. 2021). Predators may be overly cautious around humans, resulting in missed opportunities for fitness-enhancing activities. For example, Suraci et al. (2019) showed that human presence induced puma (*Puma concolor*) to abandon their kill sites. Based on camera trap data, Burton et al. (2024) showed that carnivores avoided humans to a greater extent than wild animals belonging to other foraging guilds. Chronic exposure to humans may thus lead to reduced fitness for risk-averse individuals. Alternatively, predators may not be cautious enough around humans, resulting in increased direct mortality (Smith et al. 2021). The severity and direction of the response may be proportional to the extent and duration of persecution by humans. The majority of studies we found in our search focused on primary consumers, rather than predators (Appendix A), making it challenging to assess trophic effects. This gap remains an important area for future research.

Like predators, large animals may be disproportionately targeted by humans for economic reasons and as trophies. Natural predators rarely target larger and healthier animals and thus these animals are less likely to invest in anti-predator behaviours.

Like predators, large animals may be disproportionately targeted by humans for economic reasons and as trophies. Natural predators rarely target larger and healthier animals and thus these animals are less likely to invest in anti-predator behaviours. We found no evidence for a consistent effect of body size on the behavioural responses of wild animals to human interactions in our meta-analysis (Appendix E). However, when we look at the responses of individual species across studies (Figure 4), behavioural response to lethal interactions was largest for large herbivores such as deer (*Dama dama*), elk (*Cervus* elaphus), and moose (*Alces alces*) (Deer: Pecorella et al., 2016; Elk: Ciuti et al., 2012, Proffitt et al., 2009, Shannon et al., 2014, Jayakody et al., 2008; Moose: Bhardwaj et al., 2022). Finally, most species in our dataset are group-living to some degree, while only three species (predators) are considered solitary. Group living typically reduces predation risk through the dilution and many-eyes effects (Foster & Treherne 1981; Pulliam 1973). Whether grouping similarly alters responses to humans compared to natural predators remains unclear and could not be evaluated here.

## Conclusion

Here, we present an extensive synthesis of the available literature on the effect of human interactions on animal behaviour. Although our study does not bridge the gap between behaviour. and the resulting demographic consequences for populations (see Smith et al., 2024), it is important to note that the demographic effects of nonconsumptive effects (NCEs) remain contentious (Sheriff et al., 2020). Future research should more explicitly link human interactions, behavioural changes, fitness consequences, and demographic outcomes.

Our general conclusions and suggestions for future research are below:

1. Both lethal and non–lethal interactions with humans elicit changes in wild animal behaviour., but lethal interactions have a more consistent and pronounced effect.
2. There is considerable variation in the response of animals to the different types of human interactions that has yet to be disentangled. Factors such as frequency of interactions, history of hunting, and taxon identity may modulate the response of animals to human disturbance.
3. Studies that directly compare the effects of lethal and non-lethal interactions within the same species are necessary to determine whether and when animals can infer the intent of humans.
4. Future research may benefit from contrasting humans as lethal predators with natural predators in the same space and across trophic levels, given the potential for compounding effects on animals and ecosystems.

## Acknowledgements

Funding for this research was provided by the Prime Minister’s Research Fellowship to SD, with some support from the DBT-Wellcome Trust India Alliance grant (IA/I/19/2/504639) to MT. We thank Dr. Nitya Mohanty and Dr. Daniel Noble for their invaluable input during the drafting of this manuscript. We thank Prof. Douglas W. Morris, Prof. Burt Kotler, and an anonymous reviewer for constructive comments during the review process.

## Author Contributions

The study was conceptualised by SD with substantial input from MT and KS. SD conducted data collection and analysis. The first draft of the manuscript and subsequent revisions were written with equal contribution by all authors.

## Data Accessibility Statement

Data and code used in this manuscript are available at https://doi.org/10.5281/zenodo.15099679.

## Notes

https://www.github.com/cheesesnakes/superpredator-or-superscary.

### Competing Interest Statement

The authors have declared no competing interest.

### Summary of Updates

We have revised the abstract, introduction and discussion. We have now emphasised the theoretical basis of our hypothesis and results in the introduction and abstract, respectively. We have also revised the introduction to improve the flow of ideas. Finally, we have merged two sections in the discussion for better thematic cohesion.

