## Appendix F for "Are human super-predators always super-scary? A meta-analysis of wild animal behavioural responses to human interactions"

**Appendix A: Screening protocols, PRISMA flowchart and information about studies.**

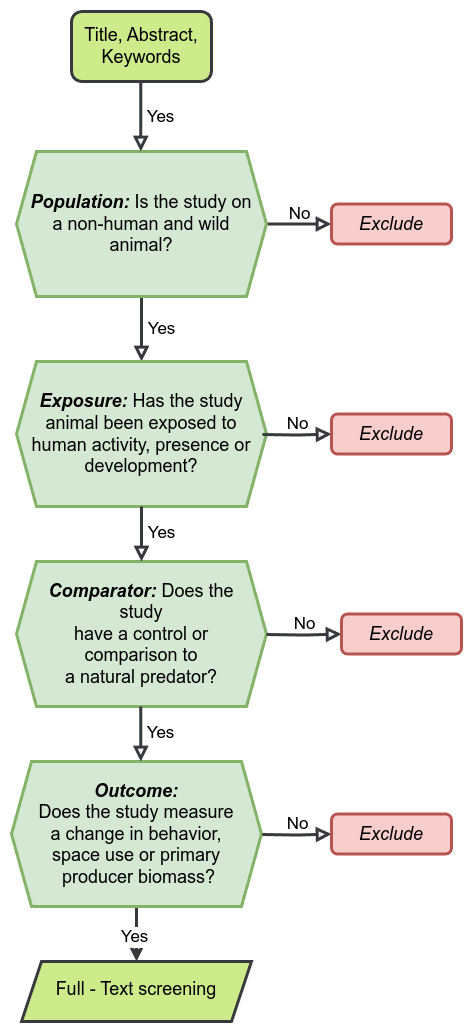

Figure A.1: Initial screening protocol.

Figure A2: Full-text screening protocol.

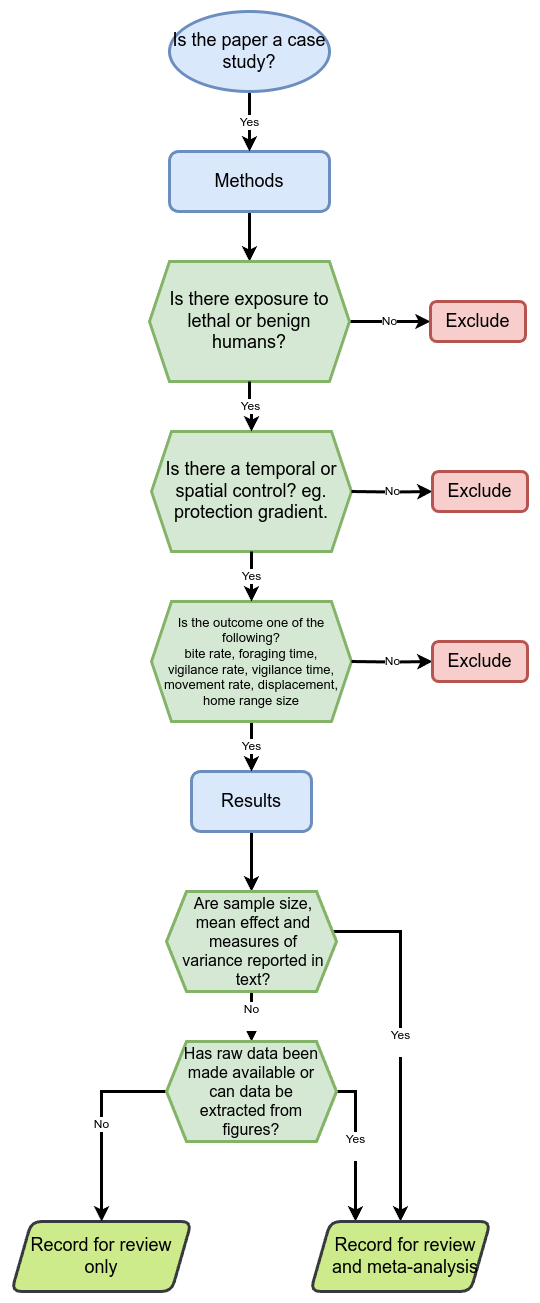

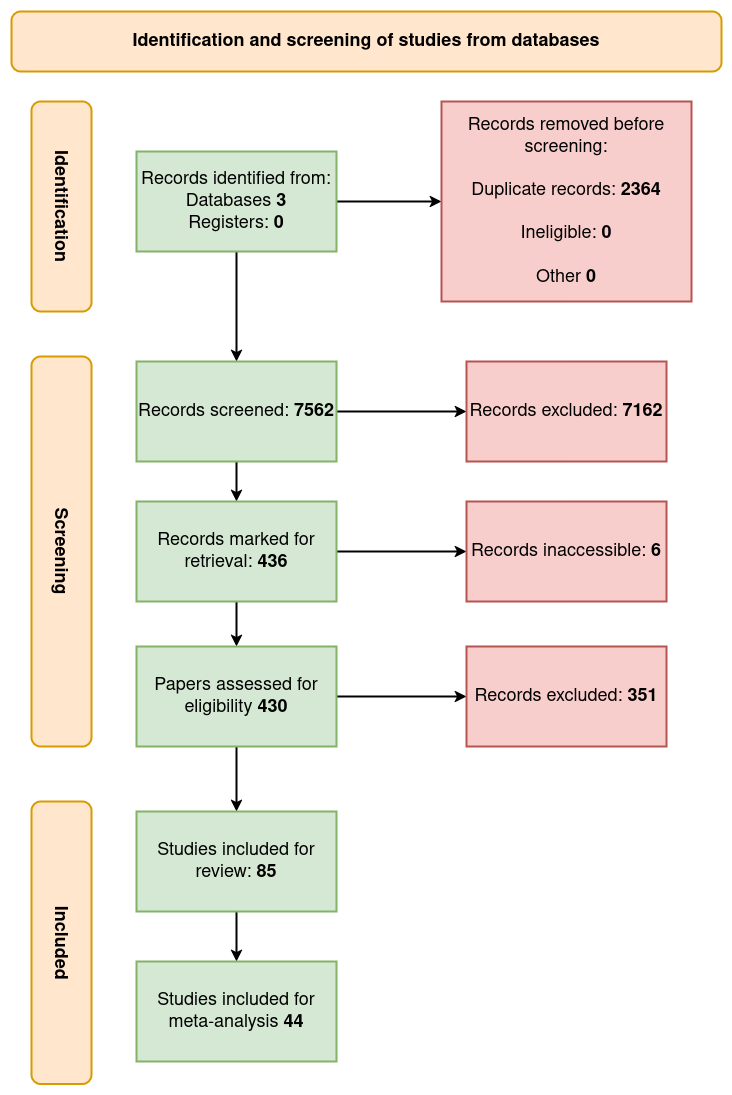

Figure A3: Summary of screening procedures. *Adapted from:*  Page MJ, McKenzie JE, Bossuyt PM, Boutron I, Hoffmann TC, Mulrow CD, et al. The PRISMA 2020 statement: an updated guideline for reporting systematic reviews. BMJ 2021;372:n71. doi: 10.1136/bmj.n71

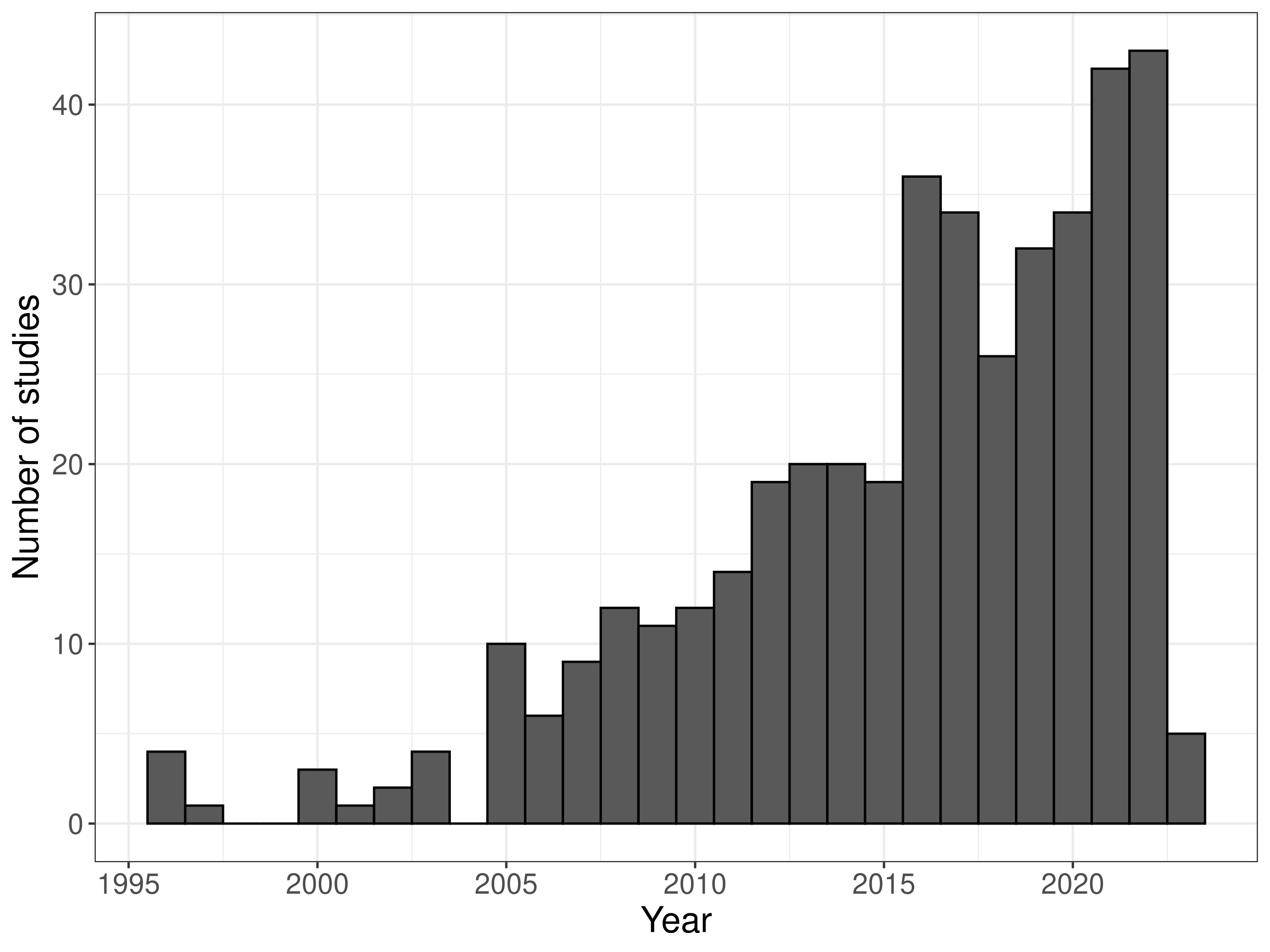

Figure A4: Number of included studies across publication years.

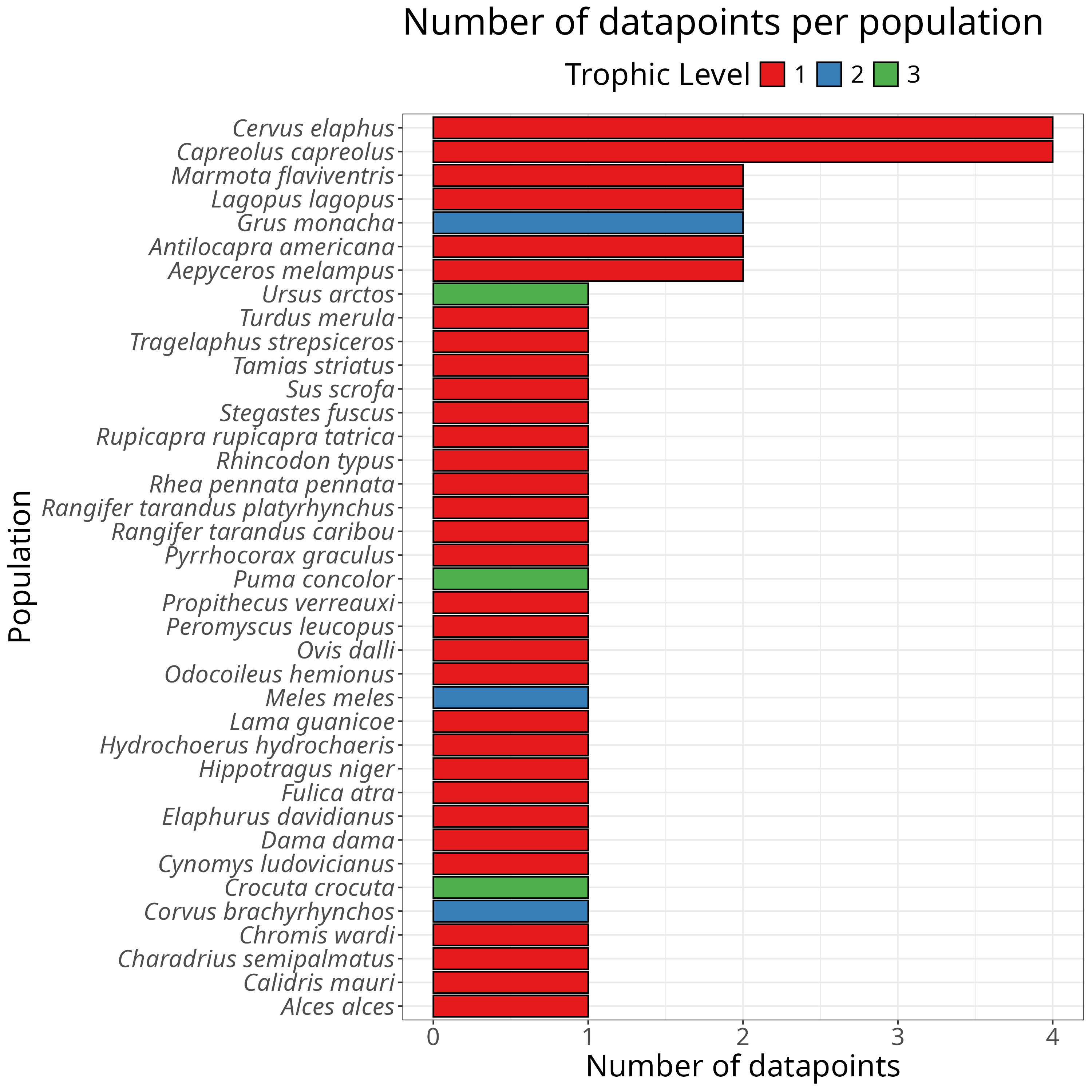

Figure A5: Number of included studies across taxa.

Table A1: Summary of study design with of studies included in the current meta-analysis including species studied, treatments used, type of contrast employed and outcome metrics measured. We have also included the exposure and outcome categories we assigned to each study as well as the trophic level and functional group of each species.

| **Citations** | **Common Name** | **Specific Epithet** | **Trophic Level** | **Functional Group** | **Contrast Type** | **Exposure Category** | **Treatment** | **Outcome**  **Category** | **Metrics** |
| --- | --- | --- | --- | --- | --- | --- | --- | --- | --- |
| Barri et al., 2012 | Lesser rhea | *Rhea pennata pennata* | 1 | Herbivore | Spatial | Lethal  interaction | Hunted | Vigilance, foraging | Foraging time , vigilance time |
| Benevides et al., 2019 | Dusky damselfish | *Stegastes fuscus* | 1 | Herbivore | Treatment–control | Active  interaction | Active disturbance | Foraging | Foraging time , habitat selection |
| Benhaiem et al., 2008 | Roe deer | *Capreolus capreolus* | 1 | Herbivore | Temporal | Lethal  interaction | Treatment | Vigilance | Feeding time , vigilance time |
| Bhardwaj et al., 2022 | Moose | *Alces alces* | 1 | Herbivore | Auditory cues | Lethal  interaction | Natural predator, active disturbance | Vigilance, foraging | Flight probability, feeding time, vigilance time |
| Broseth et al., 2010 | Willow grouse | *Lagopus lagopus* | 1 | Granivore | Spatial | Lethal  interaction | Hunting | Displacement, home range | Displacement , range size |
| Brown et al., 2020 | Mule deer | *Odocoileus hemionus* | 1 | Herbivore | Temporal | Lethal  interaction | Hunting | Displacement, home range | Movement rate , range size |
| Cappa et al., 2017 | Guanaco | *Lama guanicoe* | 1 | Herbivore | None | Passive  interaction | Passive disturbance | Vigilance, foraging | Vigilance rate pop. |
| Chen-kraus et al., 2022 | Verreaux’s sifaka | *Propithecus verreauxi* | 1 | Herbivore | None | Passive  interaction | Passive disturbance | Vigilance, foraging | Foraging time , vigilance time |
| Ciuti et al., 2012 | Elk | *Cervus elaphus* | 1 | Herbivore | Spatial | Passive  interaction | Passive disturbance | Foraging | Foraging time , vigilance time |
| Clinchy et al., 2016 | Badger | *Meles meles* | 2 | Omnivore | Auditory cues | Lethal  interaction | Hunted, natural predator, positive control | Foraging, vigilance, latency | Foraging rate , vigilance rate , foraging time , vigilance time |
| Crosmary et al., 2012 | Impala, greater kudu, sable antelope | *Aepyceros melampus, tragelaphus strepsiceros, hippotragus niger* | 1 | Herbivore | Spatial | Lethal  interaction | Hunted | Vigilance | Vigilance time |
| Olsson et al., 1996 | Willow grouse | *Lagopus lagopus* | 1 | Granivore | Spatial | Lethal  interaction | Hunting | Displacement, movement rate | Movement rate , displacement |
| Fernandez-juricic et al., 2000 | Blackbird | *Turdus merula* | 1 | Omnivore | Temporal | Active  interaction | Active disturbance | Vigilance, foraging | Vigilance time , foraging time |
| Gavin et al., 2006 | Pronghorn antelope | *Antilocapra americana* | 1 | Herbivore | Spatial | Passive  interaction | Passive disturbance | Foraging, vigilance | Vigilance time , foraging time |
| Giordano et al., 2022 | Chipmunk, white-footed mouse | *Tamias striatus,* | 1 | Herbivore | Auditory cues | Passive  interaction | Passive disturbance | Foraging, vigilance | Gud , foraging time , vigilance time |
| Grignolio et al., 2011 | Roe deer | *Capreolus capreolus* | 1 | Herbivore | Spatial | Lethal  interaction | Hunting | Home range | Range size |
| Mccormick et al., 2018 | Ward's damsel fish | *Chromis wardi* | 1 | Herbivore | None | Passive  interaction | Passive disturbance | Displacement | Displacement |
| Jayakody et al., 2008 | Red deer | *Cervus elaphus* | 1 | Herbivore | Treatment-control | Lethal  interaction, active  interaction | Active disturbance, hunted | Vigilance, foraging | Vigilance time , foraging time , vigilance frequency , foraging frequency |
| Lesmerises et al., 2017 | Gaspesie caribou | *Rangifer tarandus caribou* | 1 | Herbivore | Spatial | Active  interaction, passive  interaction | Passive disturbance | Foraging, vigilance | Foraging time , vigilance time |
| Li et al., 2011 | Yellowbellied marmot | *Marmota flaviventris* | 1 | Herbivore | None | Active  interaction | Active disturbance | Vigilance, foraging | Vigilance time , foraging time |
| Li et al., 2015 | Hooded crane | *Grus monacha* | 2 | Insectivore | Spatial | Active  interaction | Active disturbance | Foraging, vigilance | Vigilance time , foraging time |
| Li et al., 2016 | Hooded crane | *Grus monacha* | 2 | Insectivore | Spatial | Active  interaction | Active disturbance | Vigilance | Collective vigilance , vigilance time |
| Loehr et al., 2005 | Thinhorn sheep | *Ovis dalli* | 1 | Herbivore | Treatment-control | Active  interaction | Active disturbance | Vigilance, foraging | Foraging time , vigilance time |
| Lopes et al., 2021 | Capybaras | *Hydrochoerus hydrochaeris* | 1 | Herbivore | Spatial | Passive  interaction | Passive disturbance | Home range | Displacement , range size |
| Matson et al., 2005 | Impala | *Aepyceros melampus* | 1 | Herbivore | Spatial | Lethal  interaction | Hunted | Vigilance | Vigilance time , vigilance rate |
| Montero-quintana et al., 2020 | Whale shark | *Rhincodon typus* | 1 | Herbivore | Before-after control intervention | Active  interaction | Before treatment, after treatment | Foraging, vigilance | Foraging probability , vigilance probability |
| Murchison et al., 2016 | Western sandpiper, semipalmated plover | *Calidris mauri, charadrius semipalmatus* | 1 | Insectivore, herbivore | None | Active  interaction | Active disturbance | Foraging, vigilance | Density , vigilance time , foraging time |
| Pangle et al., 2010 | Spotted hyena | *Crocuta crocuta* | 3 | Carnivore | Treatment-control | Lethal  interaction, active  interaction | Hunting | Vigilance | Vigilance time |
| Pecorella et al., 2016 | Fallow deer | *Dama dama* | 1 | Herbivore | Spatial | Lethal  interaction | Hunting | Foraging, vigilance | Foraging time , vigilance time , fecal cortisol |
| Peksa et al., 2018 | Tatra chamois | *Rupicapra rupicapra tatrica* | 1 | Herbivore | None | Passive  interaction | Passive disturbance | Vigilance | Foraging time |
| Picardi et al., 2019 | Roe deer | *Capreolus capreolus* | 1 | Herbivore | Spatial | Lethal  interaction | Hunting | Movement rate | Movement rate , range size |
| Podgorski et al., 2013 | Wild boar | *Sus scrofa* | 1 | Herbivore | Spatial | Passive  interaction | Active disturbance | Displacement, home range | Displacement , range size |
| Proffitt et al., 2009 | Elk | *Cervus elaphus* | 1 | Herbivore | Temporal | Lethal  interaction | Hunting | Displacement | Movement rate , group size |
| Reimers et al., 2011 | Svalbard reindeer | *Rangifer tarandus platyrhynchus* | 1 | Herbivore | Spatial | Active  interaction | Active disturbance, hunting | Vigilance | Vigilance rate , vigilance time , fid , ad , ed |
| Rode et al., 2006 | Brown bear | *Ursus arctos* | 3 | Omnivore | Before-after control intervention | Active  interaction | Active disturbance | Foraging | Foraging time |
| Severcan et al., 2011 | Eurasian coot | *Fulica atra* | 1 | Herbivore | None | Active  interaction | Active disturbance, treatment | Vigilance, foraging | Vigilance rate , vigilance time , feeding rate |
| Shannon et al., 2014 | Pronghorn antelope, elk, prairie dog | *Antilocapra americana, cervus elaphus, cynomys ludovicianus* | 1 | Herbivore | None | Passive  interaction | Passive disturbance | Foraging, vigilance | Group size , vigilance , foraging, abundance , vigilance , foraging |
| Smith et al., 2017 | Puma | *Puma concolor* | 3 | Carnivore | Auditory cues | Lethal  interaction | Hunting | Foraging | Feeding time |
| Sonnichsen et al., 2013 | Roe deer | *Capreolus capreolus* | 1 | Herbivore | Treatment-control | Lethal  interaction | Hunting | Vigilance | Vigilance time , foraging time |
| Uchida et al., 2021 | Yellowbellied marmot | *Marmota flaviventris* | 1 | Herbivore | None | Active  interaction | Active disturbance | Vigilance | Vigilance time |
| Vallino et al., 2019 | Alpine chough | *Pyrrhocorax graculus* | 1 | Granivore | None | Active  interaction | Active disturbance | Foraging | Foraging time , prob. Flight , density |
| Ward et al., 1997 | American crow | *Corvus brachyrhynchos* | 2 | Omnivore | Spatial | Active  interaction | Passive disturbance | Vigilance | Vigilance time , foraging time |
| Zheng et al., 2013 | Pere david’s deer | *Elaphurus davidianus* | 1 | Herbivore | Spatial | Active  interaction | Active disturbance | Vigilance | Vigilance time |

Table A2: Number of studies with significant and non-significant results across types of interactions and outcomes.

| **Outcome** | **Significance** | **Direction of response** | **Active Interaction** | **Lethal Interaction** | **Passive Interaction** |
| --- | --- | --- | --- | --- | --- |
| Foraging | Non-significant | Negative | 6 | 3 | 3 |
|  |  | Positive | 4 | 0 | 0 |
|  | Significant | Negative | 4 | 3 | 3 |
|  |  | Positive | 7 | 0 | 4 |
| Movement | Non-significant | Negative | 0 | 0 | 1 |
|  |  | Positive | 0 | 6 | 1 |
|  | Significant | Negative | 0 | 3 | 2 |
|  |  | Positive | 0 | 4 | 1 |
| Vigilance | Non-significant | Negative | 4 | 1 | 0 |
|  |  | Positive | 5 | 2 | 0 |
|  | Significant | Negative | 0 | 2 | 5 |
|  |  | Positive | 16 | 8 | 2 |

Table A3: Summary of effects (SMD, 95% CI) across studies that measured multiple behavioural outcomes simultaneously.

| **Citation** | **Exposure Category** | **Outcome** | **Common Name** | **SMD** | **Upper** | **Lower** |
| --- | --- | --- | --- | --- | --- | --- |
| Barri et al., 2012 | Lethal Interaction | Foraging | Lesser Rhea | -0.453 | 0.038 | -0.944 |
|  | Lethal Interaction | Vigilance | Lesser Rhea | 0.694 | 1.193 | 0.195 |
| Bhardwaj et al., 2022 | Lethal Interaction | Foraging | Moose | -3.775 | -3.439 | -4.110 |
|  | Lethal Interaction | Vigilance | Moose | 2.073 | 2.326 | 1.820 |
| Cappa et al., 2017 | Passive Interaction | Foraging | Guanaco | 2.111 | 2.368 | 1.855 |
|  | Passive Interaction | Vigilance | Guanaco | -3.465 | -3.131 | -3.799 |
| Chen-Kraus et al., 2022 | Passive Interaction | Foraging | Verreaux’s Sifaka | 1.184 | 1.605 | 0.764 |
|  | Passive Interaction | Vigilance | Verreaux’s Sifaka | -2.141 | -1.654 | -2.628 |
| Clinchy et al., 2016 | Lethal Interaction | Foraging | Badger | -0.130 | 0.266 | -0.527 |
|  | Lethal Interaction | Vigilance | Badger | -0.079 | 0.317 | -0.475 |
| Fernandez-Juricic et al., 2000 | Active Interaction | Foraging | Blackbird | -0.305 | 0.318 | -0.928 |
|  | Active Interaction |  | Blackbird | -0.305 | 0.319 | -0.928 |
|  | Active Interaction |  | Blackbird | -0.327 | 0.297 | -0.951 |
|  | Active Interaction | Vigilance | Blackbird | 0.627 | 1.261 | -0.008 |
|  | Active Interaction |  | Blackbird | 0.680 | 1.317 | 0.042 |
|  | Active Interaction |  | Blackbird | 0.786 | 1.429 | 0.143 |
| Gavin et al., 2006 | Passive Interaction | Foraging | Pronghorn Antelope | -0.220 | 0.031 | -0.472 |
|  | Passive Interaction | Vigilance | Pronghorn Antelope | 0.283 | 0.535 | 0.031 |
| Giordano et al., 2022 | Passive Interaction | Foraging | Chipmunk | -0.385 | 0.084 | -0.855 |
|  | Passive Interaction |  | White-Footed Mouse | 1.254 | 1.828 | 0.680 |
|  | Passive Interaction | Vigilance | Chipmunk | 0.873 | 1.360 | 0.386 |
| Jayakody et al., 2008 | Active Interaction | Foraging | Elk | -0.338 | -0.053 | -0.623 |
|  | Active Interaction | Vigilance | Elk | 1.830 | 2.167 | 1.493 |
|  | Lethal Interaction | Foraging | Elk | -0.796 | -0.226 | -1.367 |
|  | Lethal Interaction | Vigilance | Elk | 6.266 | 7.266 | 5.266 |
| Lesmerises et al., 2017 | Passive Interaction | Foraging | Gaspesie Caribou | -0.268 | 0.457 | -0.993 |
|  | Passive Interaction | Vigilance | Gaspesie Caribou | -0.796 | -0.056 | -1.536 |
| Li et al., 2011 | Active Interaction | Foraging | Yellowbellied Marmot | -0.923 | -0.625 | -1.220 |
|  | Active Interaction | Vigilance | Yellowbellied Marmot | 1.550 | 1.872 | 1.227 |
| Li et al., 2015 | Active Interaction | Foraging | Hooded Crane | 0.458 | 1.101 | -0.186 |
|  | Active Interaction |  | Hooded Crane | 0.805 | 1.658 | -0.048 |
|  | Active Interaction |  | Hooded Crane | 1.414 | 1.805 | 1.022 |
|  | Active Interaction |  | Hooded Crane | 1.564 | 1.800 | 1.328 |
|  | Active Interaction | Vigilance | Hooded Crane | 0.166 | 0.519 | -0.188 |
|  | Active Interaction |  | Hooded Crane | -0.212 | 0.426 | -0.850 |
|  | Active Interaction |  | Hooded Crane | 0.367 | 0.577 | 0.156 |
|  | Active Interaction |  | Hooded Crane | -0.669 | 0.177 | -1.515 |
| Loehr et al., 2005 | Active Interaction | Foraging | Thinhorn Sheep | 0.050 | 0.670 | -0.570 |
|  | Active Interaction |  | Thinhorn Sheep | 0.437 | 1.161 | -0.287 |
|  | Active Interaction | Vigilance | Thinhorn Sheep | -0.057 | 0.659 | -0.773 |
|  | Active Interaction |  | Thinhorn Sheep | 0.944 | 1.598 | 0.291 |
| Montero-Quintana et al., 2020 | Active Interaction | Foraging | Whale Shark | -0.229 | 0.108 | -0.566 |
|  | Active Interaction | Vigilance | Whale Shark | 0.397 | 0.819 | -0.025 |
| Murchison et al., 2016 | Active Interaction | Foraging | Semipalmated Plover | -0.137 | 0.706 | -0.979 |
|  | Active Interaction |  | Western Sandpiper | -0.198 | 0.495 | -0.890 |
|  | Active Interaction | Vigilance | Semipalmated Plover | 0.495 | 1.346 | -0.356 |
|  | Active Interaction |  | Western Sandpiper | -0.191 | 0.501 | -0.883 |
| Pecorella et al., 2016 | Lethal Interaction | Foraging | Fallow Deer | -1.418 | -1.253 | -1.584 |
|  | Lethal Interaction | Vigilance | Fallow Deer | -0.768 | -0.615 | -0.922 |
| Severcan et al., 2011 | Active Interaction | Foraging | Eurasian Coot | 0.336 | 0.669 | 0.002 |
|  | Active Interaction | Vigilance | Eurasian Coot | 0.755 | 1.098 | 0.413 |
|  | Active Interaction |  | Eurasian Coot | 1.270 | 1.632 | 0.907 |
| Shannon et al. 2014 | Passive Interaction |  | Prairie Dog | 2.180 | 2.963 | 1.398 |
|  | Passive Interaction | Vigilance | Prairie Dog | -1.998 | -1.239 | -2.756 |

**Appendix B: List of studies included in the systematic review and meta-analysis. † indicates studies that were used as key studies for backward and forward search, ¥ indicates studies that were included after backward and forward search, and ^ indicates studies included in the meta-analysis**

Adam, M., M. Podhrázský, and P. Musil. 2016. “Effect of Start of Hunting Season on Behaviour of Greylag Geese Anser Anser.” Ardea 104 (1): 63–68. <https://doi.org/10.5253/arde.v104i1.a5>.

**¥** Barri, FR, N Roldan, JL Navarro, and MB Martella. 2012. “Effects of Group Size, Habitat and Hunting Risk on Vigilance and Foraging Behaviour in the Lesser Rhea (Rhea Pennata Pennata).” Emu 112 (1): 67–70. <https://doi.org/10.1071/MU10090>.

**¥** Benevides, LJ, GC Cardozo-Ferreira, CEL Ferreira, PHC Pereira, TK Pinto, and CLS Sampaio. 2019. “Fear-Induced Behavioural Modifications in Damselfishes Can Be Diver-Triggered.” Journal of Experimental Marine Biology and Ecology 514: 34–40. <https://doi.org/10.1016/j.jembe.2019.03.009>.

**¥** Benhaiem, S., M. Delon, B. Lourtet, B. Cargnelutti, S. Aulagnier, A.J.M. Hewison, N. Morellet, and H. Verheyden. 2008. “Hunting Increases Vigilance Levels in Roe Deer and Modifies Feeding Site Selection.” Animal Behaviour 76 (3): 611–18. <https://doi.org/10.1016/j.anbehav.2008.03.012>.

**¥** Bhardwaj, M., D. Lodnert, M. Olsson, A. Winsvold, S.M. Eilertsen, P. Kjellander, and A. Seiler. 2022. “Inducing Fear Using Acoustic Stimuli—A Behavioral Experiment on Moose (Alces Alces) in Sweden.” Ecology and Evolution 12 (11). <https://doi.org/10.1002/ece3.9492>.

Blackwell, BF, TL DeVault, E Fernandez-Juricic, EM Gese, L Gilbert-Norton, and SW Breck. 2016. “No Single Solution: Application of Behavioural Principles in Mitigating Human-Wildlife Conflict.” Animal Behavior 120: 245–54. <https://doi.org/10.1016/j.anbehav.2016.07.013>.

Blanc, R, M Guillemain, J Mouronval, D Desmonts, and H Fritz. 2006. “Effects of Non-Consumptive Leisure Disturbance to Wildlife.” Revue d’Écologie (La Terre et La Vie) 61 (2): 117–33. <https://doi.org/10.3406/revec.2006.1306>.

**¥** Broseth, H, and HC Pedersen. 2010. “Disturbance Effects of Hunting Activity in a Willow Ptarmigan Lagopus Lagopus Population.” Wildlife Biology 16 (3): 241–48. <https://doi.org/10.2981/09-096>.

Brown, CL, AR Hardy, JR Barber, KM Fristrup, KR Crooks, and LM Angeloni. 2012. “The Effect of Human Activities and Their Associated Noise on Ungulate Behavior.” PLOS ONE 7 (7). <https://doi.org/10.1371/journal.pone.0040505>.

**¥** Brown, CL, JB Smith, MJ Wisdom, MM Rowland, DB Spitz, and DA Clark. 2020. “Evaluating Indirect Effects of Hunting on Mule Deer Spatial Behavior.” Journal of Wildlife Management 84 (7): 1246–55. <https://doi.org/10.1002/jwmg.21916>.

**¥** Cappa, FM, SM Giannoni, and CE Borghi. 2017. “Effects of Roads on the Behaviour of the Largest South American Artiodactyl (Lama Guanicoe) in an Argentine Reserve.” Animal Behavior 131: 131–36. <https://doi.org/10.1016/j.anbehav.2017.07.020>.

**¥** Chen-Kraus, C., Raharinoro, N. A., Randrianirinarisoa, M. A., Anderson, D. J., Lawler, R. R., Watts, D. P., & Richard, A. F. (2022). Human-Lemur Coexistence in a Multiple-Use Landscape. *Frontiers in Ecology and Evolution*, *10*. <https://doi.org/10.3389/fevo.2022.779861>

**†** Chitwood, MC, C Baruzzi, and MA Lashley. 2022. “‘Ecology of Fear’ in Ungulates: Opportunities for Improving Conservation.” Ecology and Evolution 12 (3). <https://doi.org/10.1002/ece3.8657>.

**¥** Ciuti, S, JM Northrup, TB Muhly, S Simi, M Musiani, JA Pitt, and MS Boyce. 2012. “Effects of Humans on Behaviour of Wildlife Exceed Those of Natural Predators in a Landscape of Fear.” PLOS ONE 7 (11). <https://doi.org/10.1371/journal.pone.0050611>.

**¥** Clinchy, M, LY Zanette, D Roberts, JP Suraci, CD Buesching, C Newman, and DW Macdonald. 2016. “Fear of the Human ‘Super Predator’ Far Exceeds the Fear of Large Carnivores in a Model Mesocarnivore.” BEHAVIORAL ECOLOGY 27 (6): 1826–32.

**†** Crawford, Daniel A., L. Mike Conner, Michael Clinchy, Liana Y. Zanette, and Michael J. Cherry. 2022. “Prey Tells, Large Herbivores Fear the Human ‘Super Predator.’” Oecologia 198 (1): 91–98. <https://doi.org/10.1007/s00442-021-05080-w>.

Cromsigt, J P.G.M., Dries P.J. Kuijper, Marius Adam, Robert L. Beschta, Marcin Churski, Amy Eycott, Graham I.H. Kerley, Atle Mysterud, Krzysztof Schmidt, and Kate West. 2013. “Hunting for Fear: Innovating Management of Human–Wildlife Conflicts.” Edited by Jacqueline Frair. Journal of Applied Ecology 50 (3): 544–49. <https://doi.org/10.1111/1365-2664.12076>.

**¥** Crosmary, WG, P Makumbe, SD Cote, and H Fritz. 2012. “Vulnerability to Predation and Water Constraints Limit Behavioural Adjustments of Ungulates in Response to Hunting Risk.” Animal Behavior 83 (6): 1367–76. <https://doi.org/10.1016/j.anbehav.2012.03.004>.

Darimont, Chris T., Caroline H. Fox, Heather M. Bryan, and Thomas E. Reimchen. 2015. “The Unique Ecology of Human Predators.” Science 349 (6250): 858–60. <https://doi.org/10.1126/science.aac4249>.

**†** Fardell, LL, CR Pavey, and CR Dickman. 2020. “Fear and Stressing in Predator Prey Ecology: Considering the Twin Stressors of Predators and People on Mammals.” PEERJ 8. <https://doi.org/10.7717/peerj.9104>.

**† ¥** Fernández‐Juricic, E, Clavijo, M, Jiménez, MD, Asensio, E, and Lucas, E. 2001. “Bird Tolerance to Human Disturbance in Urban Parks of Madrid (Spain): Management Implications.” <https://doi.org/10.1007/978-1-4615-1531-9_12>.

**†** Frid, A, and L Dill. 2002. “Human-Caused Disturbance Stimuli as a Form of Predation Risk.” CONSERVATION ECOLOGY 6 (1). <https://doi.org/10.5751/ES-00404-060111>.

**¥** Gavin, SD, and PE Komers. 2006. “Do Pronghorn (Antilocapra Americana) Perceive Roads as a Predation Risk?” Canadian Journal of Zoology 84 (12): 1775–80. <https://doi.org/10.1139/z06-175>.

Gaynor, KM, A McInturff, and JS Brashares. 2021. “Contrasting Patterns of Risk from Human and Non-Human Predators Shape Temporal Activity of Prey.” Journal of Animal Ecology. <https://doi.org/10.1111/1365-2656.13621>.

**†** Gaynor, KM, MJ Cherry, SL Gilbert, MT Kohl, CL Larson, TM Newsome, LR Prugh, JP Suraci, JK Young, and JA Smith. 2021. “An Applied Ecology of Fear Framework: Linking Theory to Conservation Practice.” Animal Conservation 24 (3): 308–21. <https://doi.org/10.1111/acv.12629>.

Gert E.O, Willebrand, T, and Smith, A. 1996. “The Effects of Hunting on Willow Grouse *Lagopus Lagopus* Movements.” Wildlife Biology. <https://doi.org/10.2981/wlb.1996.003>.

**¥** Giordano, A., Hunninck, L., & Sheriff, M. (2022). Prey responses to predation risk under chronic road noise. *JOURNAL OF ZOOLOGY*. <https://doi.org/10.1111/jzo.12968>

**¥** Grignolio, S., E. Merli, P. Bongi, S. Ciuti, and M. Apollonio. 2011. “Effects of Hunting with Hounds on a Non-Target Species Living on the Edge of a Protected Area.” Biological Conservation 144 (1): 641–49. <https://doi.org/10.1016/j.biocon.2010.10.022>.

Harris, CM, L Thomas, EA Falcone, J Hildebrand, D Houser, PH Kvadsheim, FPA Lam, et al. 2018. “Marine Mammals and Sonar: Dose-Response Studies, the Risk-Disturbance Hypothesis and the Role of Exposure Context.” Journal of Applied Ecology 55 (1): 396–404. <https://doi.org/10.1111/1365-2664.12955>.

**†** Haswell, PM., J Kusak, and W. Hayward. 2017. “Large Carnivore Impacts Are Context-Dependent.” Food Webs, Challenges and opportunities for the study and conservation of large carnivores, 12 (September): 3–13. <https://doi.org/10.1016/j.fooweb.2016.02.005>.

Hojnowski, CE. 2017. “Spatial and Temporal Dynamics of Wildlife Use of a Human-Dominated Landscape.”

**¥** Jayakody, S, AM Sibbald, IJ Gordon, and X Lambin. 2008. “Red Deer Cervus Elephus Vigilance Behaviour Differs with Habitat and Type of Human Disturbance.” Wildlife Biology 14 (1): 81–91. [https://doi.org/10.2981/0909-6396(2008)14[81:RDCEVB]2.0.CO;2](https://doi.org/10.2981/0909-6396(2008)14%5B81:RDCEVB%5D2.0.CO;2).

Jennifer AG, Norris, K, and Sutherland, WJ. 2001. “Why Behavioural Responses May Not Reflect the Population Consequences of Human Disturbance.” Biological Conservation. <https://doi.org/10.1016/s0006-3207(00)00002-1>.

Jiang, TY, XM Wang, YZ Ding, ZS Liu, and ZH Wang. 2013. “Behavioral Responses of Blue Sheep (Pseudois Nayaur) to Nonlethal Human Recreational Disturbance.” Chinese Science Bulletin 58 (18): 2237–47. <https://doi.org/10.1007/s11434-013-5761-y>.

Keuling, O, and G Massei. 2021. “Does Hunting Affect the Behavior of Wild Pigs?” Human-Wildlife Interactions 15 (1): 44-55    WE-Science Citation Index Expanded (SCI-EXPANDED).

Kilgo, JC, Labisky, RF, and Fritzen, DE. 1998. “Influences of Hunting on the Behavior of White-Tailed Deer: Implications for Conservation of the Florida Panther.” Conservation Biology. <https://doi.org/10.1111/j.1523-1739.1998.97223.x>.

**†** Kuijper, DPJ, E Sahlen, B Elmhagen, S Chamaille-Jammes, H Sand, K Lone, and JPGM Cromsigt. 2016. “Paws without Claws? Ecological Effects of Large Carnivores in Anthropogenic Landscapes.” Proceedings of the Royal Society B - Biological Science 283 (1841). <https://doi.org/10.1098/rspb.2016.1625>.

**¥** Lesmerises, F, CJ Johnson, and MH St-Laurent. 2017. “Refuge or Predation Risk? Alternate Ways to Perceive Hiker Disturbance Based on Maternal State of Female Caribou.” Ecology and Evoloution 7 (3): 845–54. <https://doi.org/10.1002/ece3.2672>.

**¥** Li, C., Zhou, L., Li, H., & Jiang, Z. (2011). Effects of foraging mode and group pattern on vigilance behavior in water birds: A case study of mallard and black-winged stilt. *Belgian Journal of Zoology*, *141*.

**¥** Li, CL, LZ Zhou, L Xu, NN Zhao, and G Beauchamp. 2015. “Vigilance and Activity Time-Budget Adjustments of Wintering Hooded Cranes, Grus Monacha, in Human-Dominated Foraging Habitats.” PLOS ONE 10 (3). <https://doi.org/10.1371/journal.pone.0118928>.

**¥** Li, CL, G Beauchamp, Z Wang, and P Cui. 2016. “Collective Vigilance in the Wintering Hooded Crane: The Role of Flock Size and Anthropogenic Disturbances in a Human-Dominated Landscape.” Ethology 122 (12): 999–1008. <https://doi.org/10.1111/eth.12570>.

Lian, XM, TZ Zhang, YC Cao, JP Su, and S Thirgood. 2011. “Road Proximity and Traffic Flow Perceived as Potential Predation Risks: Evidence from the Tibetan Antelope in the Kekexili National Nature Reserve, China.” Wildlife Research 38 (2): 141–46. <https://doi.org/10.1071/WR10158>.

**¥** Loehr, J, M Kovanen, J Carey, H Hogmander, C Jurasz, S Karkkainen, J Suhonen, and H Ylonen. 2005. “Gender- and Age-Class-Specific Reactions to Human Disturbance in a Sexually Dimorphic Ungulate.” Canadian Journal of Zoology 83 (12): 1602–7. <https://doi.org/10.1139/Z05-162>.

**¥** Lopes, B, JF McEvoy, RG Morato, HR Luz, FB Costa, HR Benatti, TD Dias, et al. 2021. “Human-Modified Landscapes Alter Home Range and Movement Patterns of Capybaras.” Journal of Mammology 102 (1): 319–32. <https://doi.org/10.1093/jmammal/gyaa144>.

Magige, FJ, T Holmern, S Stokke, C Mlingwa, and E Røskaft. 2009. “Does Illegal Hunting Affect Density and Behaviour of African Grassland Birds? A Case Study on Ostrich (Struthio Camelus).” Biodiversity and Conservation. <https://doi.org/10.1007/s10531-008-9481-6>.

Manor, R, and D Saltz. 2003. “Impact of Human Nuisance Disturbance on Vigilance and Group Size of a Social Ungulate.” Ecological Applications 13 (6): 1830–34. <https://doi.org/10.1890/01-5354>.

**¥** Matson, T., Goldizen, A., & Putland, D. (2005). Factors affecting the vigilance and flight behaviour of impalas. *SOUTH AFRICAN JOURNAL OF WILDLIFE RESEARCH*, *35*(1), 1-11    WE-Science Citation Index Expanded (SCI-EXPANDED).

**¥** McCormick, MI., BJM Allan, H Harding, and SD Simpson. 2018. “Boat Noise Impacts Risk Assessment in a Coral Reef Fish but Effects Depend on Engine Type.” Scientific Reports 8 (1): 3847. <https://doi.org/10.1038/s41598-018-22104-3>.

Metcalfe, CA, AY Yaicurima, and S Papworth. 2022. “Observer Effects in a Remote Population of Large-Headed Capuchins, Sapajus Macrocephalus.” International Journal of Primatology 43 (2): 216–34. <https://doi.org/10.1007/s10764-021-00264-w>.

Miller, PJO, S Isojunno, E Siegal, FPA Lam, PH Kvadsheim, and C Cure. 2022. “Behavioral Responses to Predatory Sounds Predict Sensitivity of Cetaceans to Anthropogenic Noise within a Soundscape of Fear.” Proceedings of the National Academy of the United States of America 119 (13).

Moller, AP. 2012. “Urban Areas as Refuges from Predators and Flight Distance of Prey.” Behavioral Ecology 23 (5): 1030–35. <https://doi.org/10.1093/beheco/ars067>.

**¥** Montero-Quintana, AN, JA Vazquez-Haikin, T Merkling, P Blanchard, and M Osorio-Beristain. 2020. “Ecotourism Impacts on the Behaviour of Whale Sharks: An Experimental Approach.” ORYX 54 (2): 270–75. <https://doi.org/10.1017/S0030605318000017>.

Montgomery, RA, DW Macdonald, and MW Hayward. 2020. “The Inducible Defences of Large Mammals to Human Lethality.” Funtional Ecology 34 (12): 2426–41. <https://doi.org/10.1111/1365-2435.13685>.

Montgomery, RA., J Raupp, SAMiller, M Wijers, R Lisowsky, A Comar, CK. Bugir, and MW. Hayward. 2022. “The Hunting Modes of Human Predation and Potential Nonconsumptive Effects on Animal Populations.” Biological Conservation 265 (January): 109398. <https://doi.org/10.1016/j.biocon.2021.109398>.

**¥** Murchison, CR, Y Zharikov, and E Nol. 2016. “Human Activity and Habitat Characteristics Influence Shorebird Habitat Use and Behavior at a Vancouver Island Migratory Stopover Site.” Environmental Management 58 (3): 386–98. <https://doi.org/10.1007/s00267-016-0727-x>.

Newey, P. 2007. “Foraging Behaviour of the Common Myna (Acridotheres Tristis) in Relation to Vigilance and Group Size.” Emu 107 (4): 315–20. <https://doi.org/10.1071/MU06054>.

**¥** Olsson GE, Willebrand T, Smith, A. . (1996). The effects of hunting on willow grouse Lagopus lagopus movements. *Wildlife Biology*. <https://doi.org/10.2981/wlb.1996.003>

**†** Ordiz, A, R Bischof, and JE Swenson. 2013. “Saving Large Carnivores, but Losing the Apex Predator?” Bilogical Conservation 168: 128–33. <https://doi.org/10.1016/j.biocon.2013.09.024>.

Ordiz, A, M Aronsson, J Persson, OG Stoen, JE Swenson, and J Kindberg. 2021. “Effects of Human Disturbance on Terrestrial Apex Predators.” Diversity-Basel 13 (2).

**†** Oriol-Cotterill, A, M Valeix, LG Frank, C Riginos, and DW Macdonald. 2015. “Landscapes of Coexistence for Terrestrial Carnivores: The Ecological Consequences of Being Downgraded from Ultimate to Penultimate Predator by Humans.” OIKOS 124 (10): 1263–73. <https://doi.org/10.1111/oik.02224>.

Parsons, AW, C Bland, T Forrester, MC Baker-Whatton, SG Schuttler, WJ McShea, R Costello, and R Kays. 2016. “The Ecological Impact of Humans and Dogs on Wildlife in Protected Areas in Eastern North America.” Biological Conservation 203: 75–88. <https://doi.org/10.1016/j.biocon.2016.09.001>.

**¥** Pecorella, I, F Ferretti, A Sforzi, and E Macchi. 2016. “Effects of Culling on Vigilance Behaviour and Endogenous Stress Response of Female Fallow Deer.” Wildlife Research 43 (3): 189–96. <https://doi.org/10.1071/WR15118>.

**¥** Peksa, L, and M Ciach. 2018. “Daytime Activity Budget of an Alpine Ungulate (Tatra Chamois Rupicapra Rupicapra Tatrica): Influence of Herd Size, Sex, Weather and Human Disturbance.” Mammal Research 63 (4): 443–53. <https://doi.org/10.1007/s13364-018-0376-y>.

**¥** Podgorski, T, G Bas, B Jedrzejewska, L Sonnichsen, S Sniezko, W Jedrzejewski, and H Okarma. 2013. “Spatiotemporal Behavioral Plasticity of Wild Boar (Sus Scrofa) under Contrasting Conditions of Human Pressure: Primeval Forest and Metropolitan Area.” Journal of Mammology 94 (1): 109–19. <https://doi.org/10.1644/12-MAMM-A-038.1>.

**¥** Picardi, S, M Basille, W Peters, JM Ponciano, L Boitani, and F Cagnacci. 2019. “Movement Responses of Roe Deer to Hunting Risk.” Journal of Wildlife Management 83 (1): 43–51. <https://doi.org/10.1002/jwmg.21576>.

Price, M. 2008. “The Impact of Human Disturbance on Birds: A Selective Review.” Australian Zoologist 34: 163–96. <https://doi.org/10.7882/fs.2008.023>.

**¥** Proffitt, KM, JL Grigg, KL Hamlin, and RA Garrott. 2009. “Contrasting Effects of Wolves and Human Hunters on Elk Behavioral Responses to Predation Risk.” Journal of Wildlife Management 73 (3): 345–56. <https://doi.org/10.2193/2008-210>.

Proudman, NJ, M Churski, JW Bubnicki, JA Nilsson, and DPJ Kuijper. 2021. “Red Deer Allocate Vigilance Differently in Response to Spatio-Temporal Patterns of Risk from Human Hunters and Wolves.” Wildlife Research 48 (2): 163–74. <https://doi.org/10.1071/WR20059>.

**¥** Pangle, WM, and KE Holekamp. 2010. “Lethal and Nonlethal Anthropogenic Effects on Spotted Hyenas in the Masai Mara National Reserve.” Journal of Mammology 91 (1): 154–64. <https://doi.org/10.1644/08-mamm-a-359r.1>.

**¥** Reimers, E., Lund, S., & Ergon, T. (2011). Vigilance and fright behaviour in the insular Svalbard reindeer (Rangifer tarandus platyrhynchus). *CANADIAN JOURNAL OF ZOOLOGY*, *89*(8), 753–764. <https://doi.org/10.1139/Z11-040>

Reisland, MA, N Malone, and JE Lambert. 2021. “Endangered Apes-Can Their Behaviors Be Used to Index Fear and Disturbance in Anthropogenic Landscapes?” Diversity-Basel 13 (12).

**¥** Rode, KD, SD Farley, and CT Robbins. 2006. “Behavioral Responses of Brown Bears Mediate Nutritional Effects of Experimentally Introduced Tourism.” Biological Conservation 133 (1): 70–80. <https://doi.org/10.1016/j.biocon.2006.05.021>.

Rosell, F, and Czech, A. 2000. “Responses of Foraging Eurasian Beavers Castor Fiber to Predator Odours.”*Wildlife Biology* 6 (1): 13–21. <https://doi.org/10.2981/wlb.2000.033>.

**¥** Severcan, C, and E Yamac. 2011. “The Effects of Flock Size and Human Presence on Vigilance and Feeding Behavior in the Eurasian Coot (Fulica Atra L.) during Breeding Season.” Acta Ethologica 14 (1): 51–56. <https://doi.org/10.1007/s10211-010-0089-y>.

Schuttler, S. G., A. W. Parsons, T. D. Forrester, M. C. Baker, W. J. McShea, R. Costello, and R. Kays. 2017. “Deer on the Lookout: How Hunting, Hiking and Coyotes Affect White-Tailed Deer Vigilance.” Journal of Zoology 301 (4): 320–27. <https://doi.org/10.1111/jzo.12416>.

**¥** Shannon, G, LS Cordes, AR Hardy, LM Angeloni, and KR Crooks. 2014. “Behavioral Responses Associated with a Human-Mediated Predator Shelter.” PLOS ONE 9 (4). <https://doi.org/10.1371/journal.pone.0094630>.

**¥** Shannon, G, LM Angeloni, G Wittemyer, KM Fristrup, and KR Crooks. 2014. “Road Traffic Noise Modifies Behaviour of a Keystone Species.” Animal Behavior 94: 135–41. <https://doi.org/10.1016/j.anbehav.2014.06.004>.

**† ¥** Smith, JA, JP Suraci, M Clinchy, A Crawford, D Roberts, LY Zanette, and CC Wilmers. 2017. “Fear of the Human ‘super Predator’ Reduces Feeding Time in Large Carnivores.” Proceedings of Royal Society B - Biological Sciences 284 (1857). <https://doi.org/10.1098/rspb.2017.0433>.

**†** Smith, JA, KM Gaynor, and JP Suraci. 2021. “Mismatch Between Risk and Response May Amplify Lethal and Non-Lethal Effects of Humans on Wild Animal Populations.” Frontiers in Ecology and Evolution 9 (March): 604973. <https://doi.org/10.3389/fevo.2021.604973>.

**¥** Sonnichsen, L, M Bokje, J Marchal, H Hofer, B Jedrzejewska, S Kramer-Schadt, and S Ortmann. 2013. “Behavioural Responses of European Roe Deer to Temporal Variation in Predation Risk.” Ethology 119 (3): 233–43. <https://doi.org/10.1111/eth.12057>.

Stillman, RA, and JD Goss-Custard. 2002. “Seasonal Changes in the Response of Oystercatchers Haematopus Ostralegus to Human Disturbance.” Journal of Avian Biology 33 (4): 358–65. <https://doi.org/10.1034/j.1600-048X.2002.02925.x>.

**†** Storch, I. 2013. “Human Disturbance of Grouse - Why and When?” Wildlife Biology 19 (4): 390–403. <https://doi.org/10.2981/13-006>.

**†** Tablado, Z, and L Jenni. 2017. “Determinants of Uncertainty in Wildlife Responses to Human Disturbance.” Biological Reviews 92 (1): 216–33. <https://doi.org/10.1111/brv.12224>.Theuerkauf J, W Jędrzejewski, K Schmidt, and R Gula. 2003. “Spatiotemporal Segregation of Wolves from Humans in the Bialowieza Forest (POLAND).” Journal of Wildlife Management. <https://doi.org/10.2307/3802677>.

Tucker, MA., K Böhning-Gaese, WF Fagan, JM Fryxell, B Van Moorter, SC Alberts, AH Ali, et al. 2018. “Moving in the Anthropocene: Global Reductions in Terrestrial Mammalian Movements.” Science 359 (6374): 466–69. <https://doi.org/10.1126/science.aam9712>.

Tsunoda, H. 2021. “Human Disturbances Increase Vigilance Levels in Sika Deer (Cervus Nippon): A Preliminary Observation by Camera-Trapping.” Russian Journal of Theriology 20 (1): 59–69. <https://doi.org/10.15298/rusjtheriol.20.1.07>.

**¥** Uchida, K., & Blumstein, D. (2021). Habituation or sensitization? Long-term responses of yellow-bellied marmots to human disturbance. *BEHAVIORAL ECOLOGY*, *32*(4), 668–678. <https://doi.org/10.1093/beheco/arab016>

**¥** Vallino, C, E Caprio, F Genco, D Chamberlain, et al. 2019. “Behavioural Responses to Human Disturbance in an Alpine Bird.” Journal of Ornithology. <https://doi.org/10.1007/s10336-019-01660-z>.

**¥** Ward, C, and Low, B.S. 1997. “Predictors of Vigilance for American Crows Foraging in an Urban Environment.” *The Wilson Journal of Ornithology*.

Yasue, M. 2005. “The Effects of Human Presence, Flock Size and Prey Density on Shorebird Foraging Rates.” Journal of Ethology 23 (2): 199–204. <https://doi.org/10.1007/s10164-005-0152-8>.

Yasue, M, P Dearden, and A Moore. 2008. “An Approach to Assess the Potential Impacts of Human Disturbance on Wintering Tropical Shorebirds.” ORYX 42 (3): 415–23. <https://doi.org/10.1017/S0030605308061139>.

Zanette, LY, and M Clinchy. 2020. “Ecology and Neurobiology of Fear in Free-Living Wildlife.” Annual review of Ecology, Evolution and Systematics, VOL 51, 2020 51: 297–318.

**¥** Zheng, W, G Beauchamp, XL Jiang, ZQ Li, and QL Yang. 2013. “Determinants of Vigilance in a Reintroduced Population of Pere David’s Deer.” Current Zoology 59 (2): 265–70. <https://doi.org/10.1093/czoolo/59.2.265>.

**Appendix C: Selected Examples**

**Table C1:** Examples of studies that contrast human interactions and natural predators.

| **Contrast** | **Examples** |
| --- | --- |
| Lethal human interactions had a larger effect than non – lethal human effects | Jayakody et al. 2008, Pangle and Holekamp 2010, Ciuti et al. 2012 |
| Lethal human interactions had a larger effect than natural predators | Proffitt et al. 2009, Sonnichsen et al. 2013, Clinchy et al. 2016, Parsons et al. 2016, Gaynor et al. 2021 |
| No effects across lethal and non –lethal human interactions and natural predators | Schuttler et al. 2017 |

**Table C2:** Review papers that address the behavioural consequences of human interactions across taxa. See appendix B for full citations.

| **Taxa** | **Reviews** | **Type of interaction** |
| --- | --- | --- |
| Birds | Price 2008, Storch 2013 | Active lethal interactions where taxa are hunted |
| Carnivores | Ordiz et al. 2013, Oriol Cotterill et al. 2015, Haswell et al. 2017 |  |
| Carnivores | Kuijper et al. 2016 | Active non-lethal interactions |
| Mammals | Montgomery et al. 2020, Keuling et al. 2021, Zanette et al. 2020, Chitwood et al. 2022 |  |
| Marine mammals | Hariss et al. 2018 |  |
| Cross Taxa | Tablado et al. 2017 |  |

**Table C3:** Examples illustrating the behavioural consequences of human interactions on animals

| **Type of interactions** | **‍Behavior** | **Change in behavior** | **Examples** |
| --- | --- | --- | --- |
| **Lethal** | ‍Foraging | Reduced foraging behaviour in response to hunters and fishers | Rosell 2000, Benhaiem et al. 2008, Ciuti et al. 2012, Barri et al. 2012, Clinchy et al. 2016, Pecorella et al. 2016, Smith et al. 2017, Benevides et al. 2019 |
|  | ‍Vigilance | Increased vigilance in response to human hunters | Jayakody et al. 2008, Benhaiem et al. 2008, Pangle and Holekamp 2010, Crosmary et al. 2012, Ciuti et al. 2012, Barri et al. 2012, Clinchy et al. 2016, Pecorella et al. 2016, Crawford et al. 2022 |
|  | ‍Movement  ‍ | Altered movement rates in response to human hunters | Proffitt et al. 2009 |
|  |  | Reduced movement in response to human hunters | Broseth and Pedersen 2010, Grignodio et al. 2011, Podgorsky et al. 2013, Adam et al. 2016, Picardi et al. 2019, Brown et al. 2020 |
| **Active Non-lethal** | ‍Foraging | Reduced foraging in response to the presence of tourists, vehicles, etc. | Stillman 2002, Loehr 2005, Yasue 2005, Newey 2007, Yasue 2008, Brown et al. 2012, Peska et al. 2018, Montero – Quintana et al. 2020 |
|  | ‍Vigilance  ‍ | Increased vigilance in response to human presence | Manor 2003, Loehr 2005, Newey 2007; Jayakody et al. 2008, Jiang et al. 2013, Zheng et al. 2013, Li et al. 2016, Lesmerises et al. 2017, Montero – Quintana et al. 2020 |
|  |  | Altered grouping behaviour in response to human presence | Yasue 2005 |
|  | ‍ | No effect | Rode et al. 2006, Servercan and Yamac 2011, Murchison et al. 2016 |
| **Passive Non-lethal** | ‍Foraging | Reduced foraging behaviour in response to roads, house, etc. | Gavin 2006, Brown et al. 2012, Shannon et al. 2014 |
|  | ‍Vigilance | Increased vigilance in response to roads, houses, etc | Gavin 2006, Lian et al. 2011, Shannon et al. 2014, Lesmerises et al. 2017 |
|  | ‍Movement | Altered movement in response to roads | McCormick et al. 2018 |
|  | ‍ | No effect | Cappa et al. 2107 |

**Appendix D: Sensitivity and robustness of meta-analysis.**

*Sensitivity analysis*

Our analysis draws from a global cross taxonomic dataset and, thus, there are multiple possible sources of non-independence. Bias in terms of the taxa studies and geographic location of study meant that certain populations were represented multiple times in our data set. Phylogenetic relatedness may also lead to non-independence across effect sizes(Noble et al. 2017). We account for potential non-independence by implementing the MLMA with a random effect. We evaluated publication bias visually in our dataset using a funnel plot and used regression test for funnel plot asymmetry (Nakagawa et al. 2022). We also used a multilevel meta-regression on sample variance with study identity and data identity as random effect to determine publication bias. As significant results have a tendency to be published first, we plotted the effect size against the year of publication to determine any time-lag effects. To determine the robustness of our analysis, we used a leave-one-out method where we dropped each study from our data set in sequence and ran the MLMA to determine if any single study had a disproportionate effect on the summary effect size (Nakagawa et al. 2022; Noble et al. 2017).

The regression test for asymmetry suggests significant publication bias towards negative effects in studies reporting vigilance behaviours ( = -1.059 95% CI = -3.145 – 1.027, Z = 2.25, p = 0.015) and foraging behaviours ( = -1.463 95% CI = -2.562 – -0.364, Z = 2.141, p = 0.039). The regression test for asymmetry suggests significant publication bias towards positive effects in studies of animal movement ( = 1.531 95% CI = 0.37 – 2.693, Z = -3.084, p = 0.007). The regression on variance revealed a significant effect of studies with small sample size on both vigilance and movement but not on foraging. We did not find any evidence for time-lag effects nor did any single study have disproportionate effect on our overall effect size.

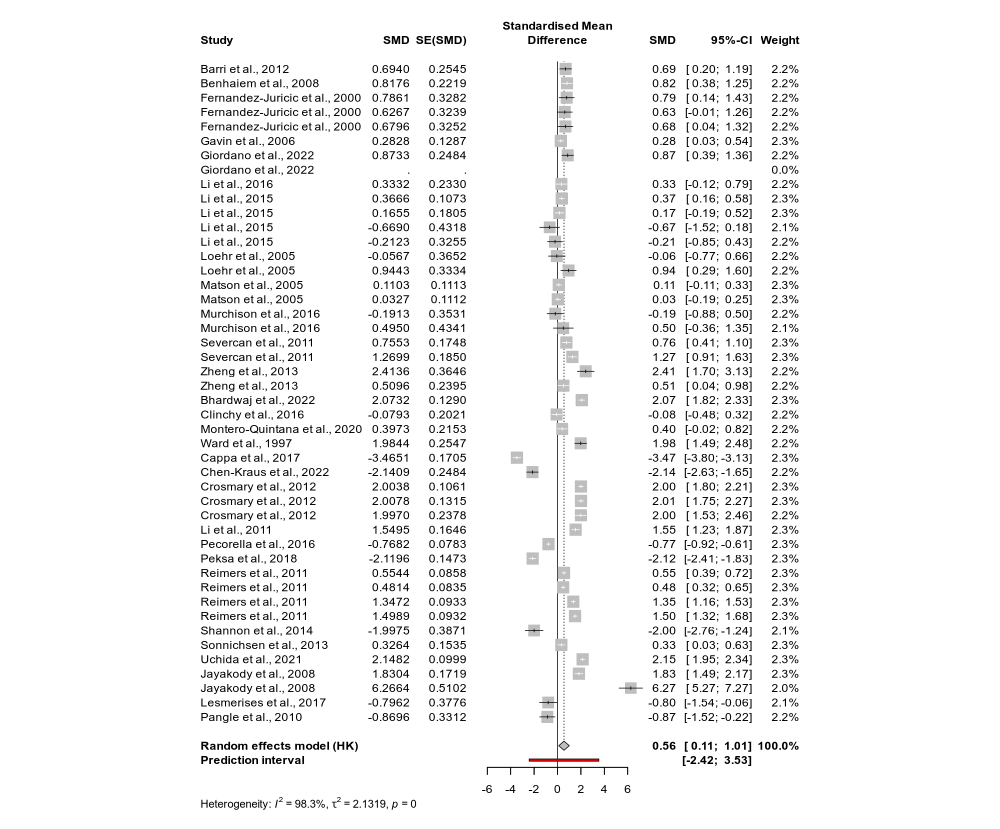

Figure D1: Forest plot of vigilance studies depicting high heterogeneity across studies ($I^{2}$ = 98.3%).

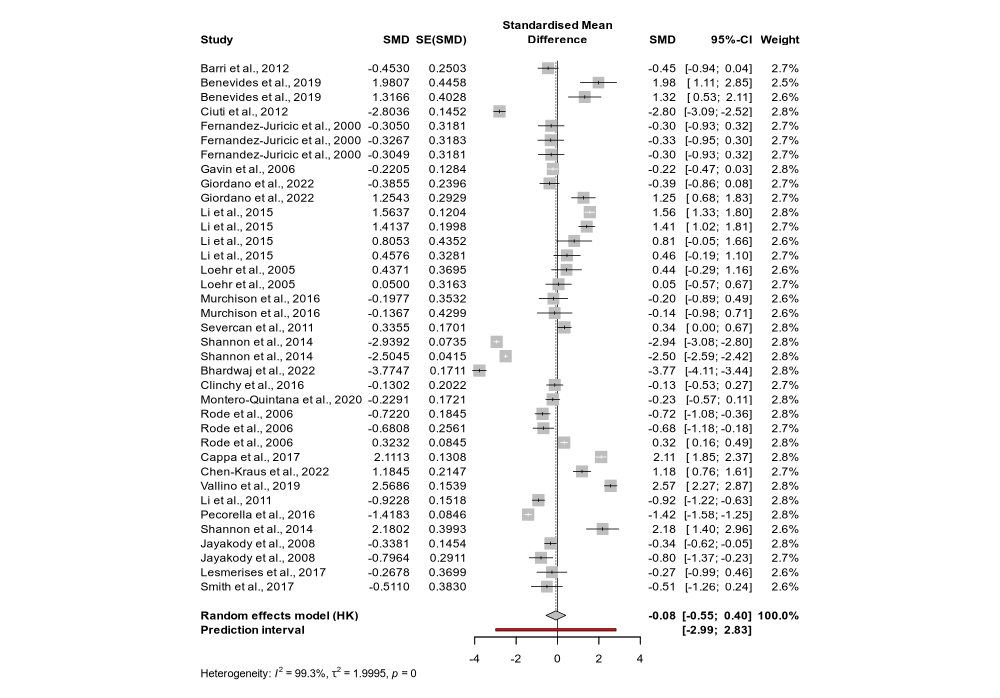

Figure D2: Forest plot of foraging studies depicting high heterogeneity across studies ($I^{2}$ = 99.3%).

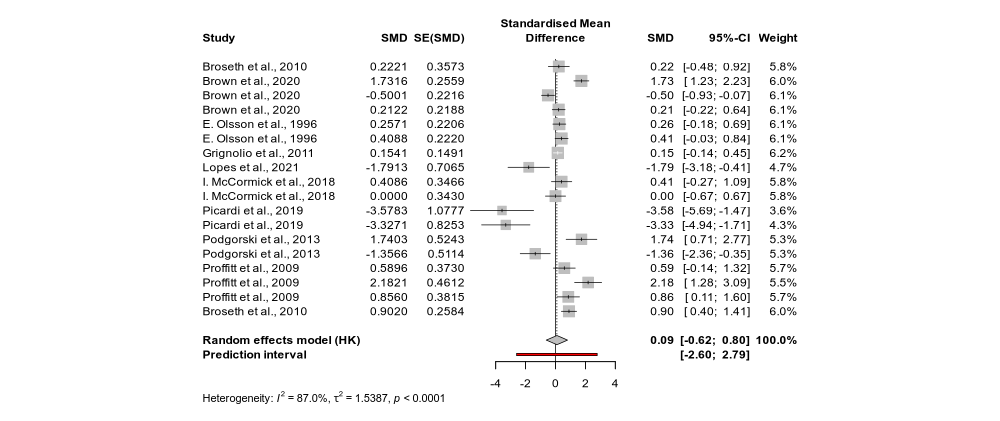

Figure D3: Forest plot of Movement studies depicting high heterogeneity across studies ($I^{2}$ = 87%).

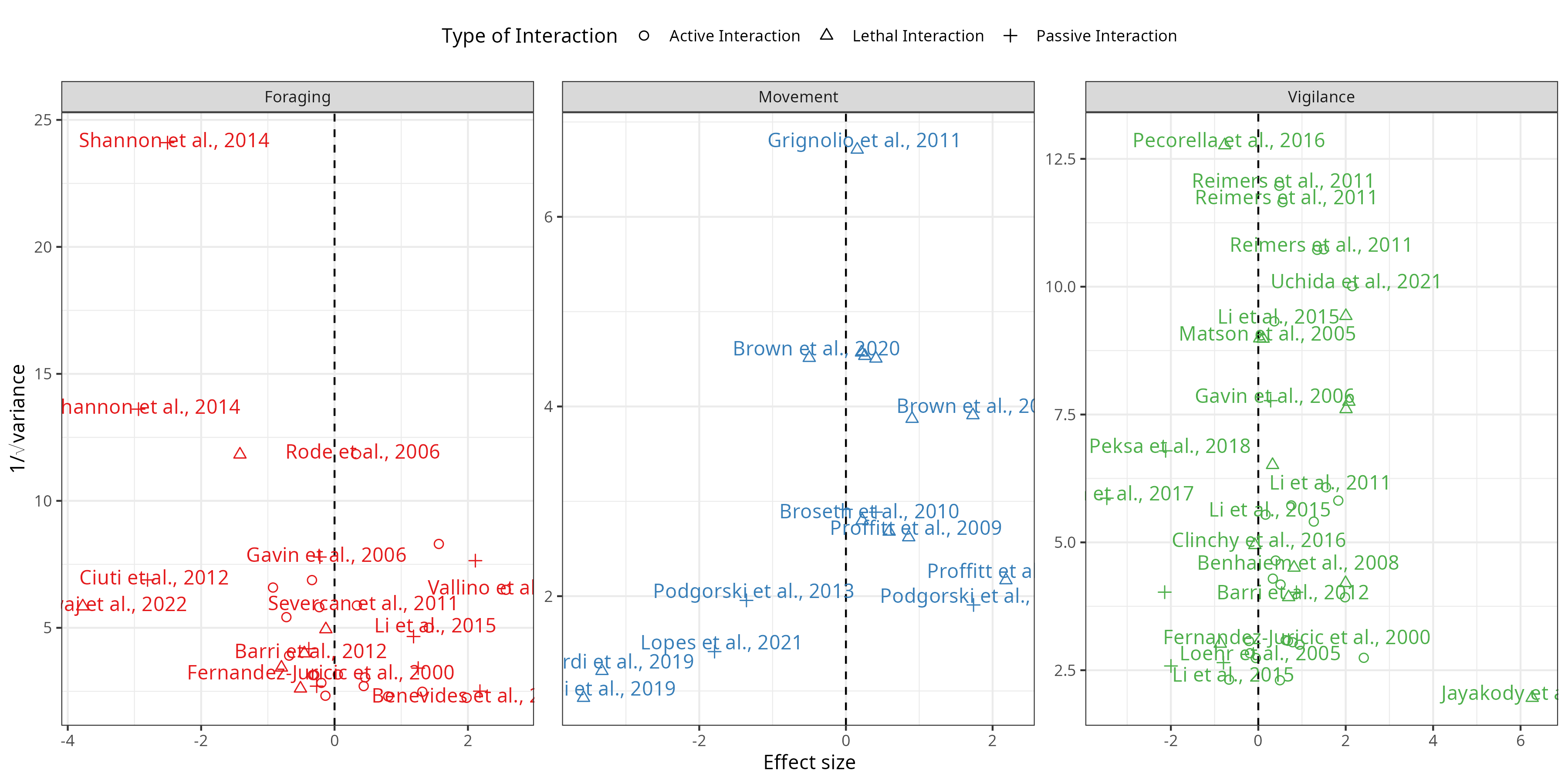

Figure D4: Funnel plot to evaluate publication bias in vigilance, foraging and movement studies.

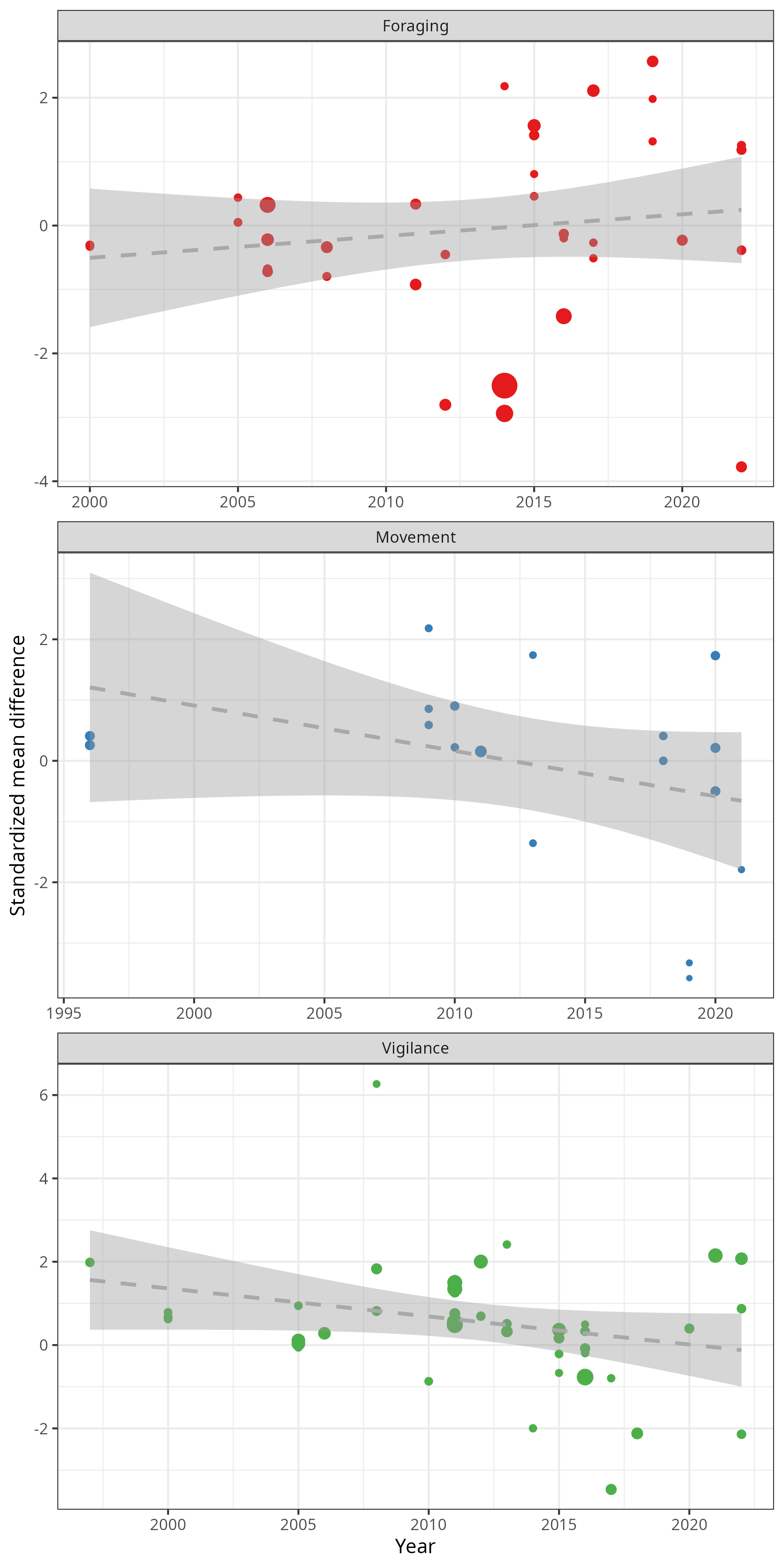

Figure D5: Plot of effect size across year of publication shows no significant correlation suggesting the lack of a time-lag effect.

Figure D6: Leave one out analysis suggests that no study had a significant impact on the final inference.

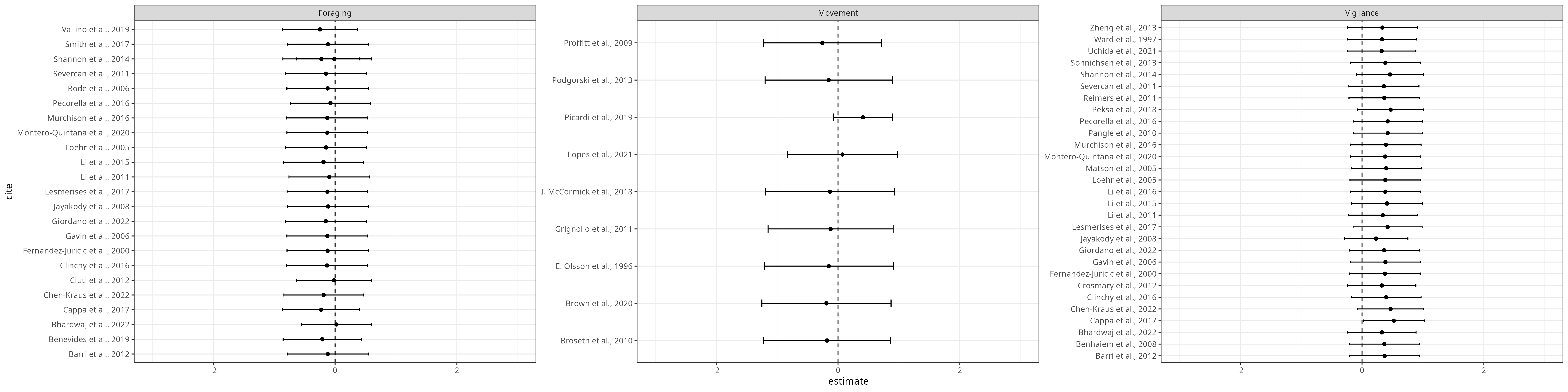

Table D1: Regression on variance to determine publication bias.

| **Term** | **Estimate** | **Standard Error** | **Statistic** | **P-value** | **Outcome** |
| --- | --- | --- | --- | --- | --- |
| Intercept | -0.04 | 0.34 | -0.11 | 0.91 | Vigilance |
| Modifiers | 6.65 | 2.99 | 2.22 | 0.03 |  |
| Intercept | -0.10 | 0.37 | -0.25 | 0.80 | Foraging |
| Modifiers | -0.55 | 3.06 | -0.18 | 0.86 |  |
| Intercept | 0.89 | 0.29 | 3.05 | 0.01 | Movement |
| Modifiers | -4.12 | 1.03 | -3.99 | 0.00 |  |

**Appendix E: Model results for meta-regression**

Table E1: Summary of model with human interactions as moderator

| Term | Estimate | Standard Error | Statistic | P-value | Outcome |
| --- | --- | --- | --- | --- | --- |
| Intercept | 0.522 | 0.356 | 1.467 | 0.155 | Vigilance |
| Size | 0.000 | 0.000 | -0.070 | 0.944 |  |
| Lethal Interaction | 0.806 | 0.496 | 1.625 | 0.111 |  |
| Passive Interaction | -1.861 | 0.619 | -3.005 | 0.006 |  |
| Intercept | 0.204 | 0.455 | 0.448 | 0.659 | Foraging |
| Size | 0.000 | 0.000 | -0.298 | 0.767 |  |
| Lethal Interaction | -0.932 | 0.515 | -1.811 | 0.079 |  |
| Passive Interaction | -0.249 | 0.709 | -0.351 | 0.729 |  |
| Intercept | -0.387 | 0.697 | -0.555 | 0.599 | Movement |
| Size | 0.006 | 0.006 | 0.948 | 0.380 |  |
| Passive Interaction | -0.228 | 1.018 | -0.224 | 0.831 |  |

Table E2: Summary of coefficients for intercept only non-phylogenetic model.

| Outcome | Intercept | SE Intercept | Sigma^2 | Tau^2 | T-value | Q-statistic | Df |
| --- | --- | --- | --- | --- | --- | --- | --- |
| Vigilance | 0.383 | 0.275 | 0.000 | 0 | 1.392 | 2,649.911 | 45 |
| Foraging | -0.097 | 0.315 | 0.363 | 0 | -0.307 | 4,920.567 | 37 |
| Movement | -0.074 | 0.428 | 0.029 | 0 | -0.172 | 130.356 | 18 |

Table E3: Random effects for intercept only non-phylogenetic model

| Outcome | Factor | Sigma^2 | N Levels |
| --- | --- | --- | --- |
| Vigilance | Species | 0.000 | 28 |
|  | Study ID | 1.636 | 29 |
|  | Data ID | 0.662 | 45 |
| Foraging | Species | 0.363 | 24 |
|  | Study ID | 1.803 | 24 |
|  | Data ID | 0.165 | 37 |
| Movement | Species | 0.029 | 7 |
|  | Study ID | 1.041 | 9 |
|  | Data ID | 0.816 | 18 |

Table E4: Summary of coefficients for intercept only phylogenetic model

| Outcome | Intercept | SE Intercept | Sigma^2 | Tau^2 | T-value | Q-statistic | Df |
| --- | --- | --- | --- | --- | --- | --- | --- |
| Vigilance | 0.364 | 0.286 | 0.000 | 0 | 1.271 | 2,479.117 | 41 |
| Foraging | 0.141 | 0.531 | 0.188 | 0 | 0.266 | 4,920.567 | 37 |
| Movement | -0.130 | 0.500 | 0.052 | 0 | -0.260 | 129.363 | 16 |

Table E5: Random effects for phylogenetic model

| Outcome | Factor | Sigma^2 | N Levels |
| --- | --- | --- | --- |
| Vigilance | Species | 0.000 | 27 |
|  | Study ID | 1.632 | 28 |
|  | Data ID | 0.780 | 41 |
|  | Phylogeny | 0.000 | 27 |
| Foraging | Species | 0.188 | 24 |
|  | Study ID | 1.415 | 24 |
|  | Data ID | 0.173 | 37 |
|  | Phylogeny | 0.761 | 24 |
| Movement | Species | 0.052 | 6 |
|  | Study ID | 1.278 | 8 |
|  | Data ID | 0.932 | 16 |
|  | Phylogeny | 0.000 | 6 |
